# Invariant neural dynamics drive commands to control different movements

**DOI:** 10.1101/2021.08.27.457931

**Authors:** Vivek R. Athalye, Preeya Khanna, Suraj Gowda, Amy L. Orsborn, Rui M. Costa, Jose M. Carmena

## Abstract

It has been proposed that the nervous system has the capacity to generate a wide variety of movements because it re-uses some invariant code. Previous work has identified that dynamics of neural population activity are similar during different movements, where dynamics refer to how the instantaneous spatial pattern of population activity changes in time. Here we test whether invariant dynamics of neural populations are actually used to issue the commands that direct movement. Using a brain-machine interface that transformed rhesus macaques’ motor cortex activity into commands for a neuroprosthetic cursor, we discovered that the same command is issued with different neural activity patterns in different movements. However, these different patterns were predictable, as we found that the transitions between activity patterns are governed by the same dynamics across movements. These invariant dynamics are low-dimensional, and critically, they align with the brain-machine interface, so that they predict the specific component of neural activity that actually issues the next command. We introduce a model of optimal feedback control that shows that invariant dynamics can help transform movement feedback into commands, reducing the input that the neural population needs to control movement. Altogether our results demonstrate that invariant dynamics drive commands to control a variety of movements, and show how feedback can be integrated with invariant dynamics to issue generalizable commands.

## Introduction

Our brain can generate a vast variety of movements. It is believed that the brain would not have such capacity if it used separate populations of neurons to control each movement. Thus, it has been proposed that the brain’s capacity to produce different movements relies on re-using the dynamics of a specific neural population’s activity ^1–3^. While theoretical work shows how dynamics emerge from neural activity transmitted through recurrent connectivity^1, 4–6^, it has been elusive to identify whether the brain re-uses dynamics to actually control movements.

Recent work on the motor cortex, a region that controls movement through direct projections to the spinal cord ^7^ and other motor centers ^8–10^, has found that population dynamics are similar across different movements. Specifically, the spatial pattern of population activity at a given time point (i.e. the instantaneous firing rate of each neuron in the population) systematically influences what spatial pattern occurs next. Models of dynamics ℎ that are invariant across movements^3^ can predict the transition from the current population activity pattern 𝑥_𝑡_ to the subsequent pattern 𝑥_𝑡+1_:

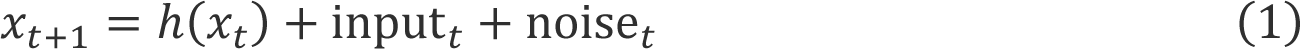

where external input input_𝑡_ and noise noise_𝑡_ are typically unmeasured. Recent work^11^ has provided the intuition that invariant dynamics bias neural activity to avoid “tangling” – which is when the same activity pattern undergoes different transitions in different movements. These dynamics models have explained features of neural activity that were unexpected from behavior ^11–14^ such as oscillations^12^, and have predicted neural activity during different movements on single trials ^15–18^, for single neurons’ spiking ^15^, for local field potential features ^19, 20^, and over many days ^18, 21^. These models also help predict behavior ^16, 18, 19, 22^.

While past work characterized the statistical relationship between invariant dynamics and behavior, it remains untested if invariant dynamics are actually used to issue commands for movement. This test requires identifying the causal transformation from neural activity to command, where the “command” is the instantaneous influence of the nervous system on movement. This is a long-standing challenge in motor control. While past work has modeled this transformation^23–25^, ongoing research reveals its complexity^8–10, 26–28^.

We addressed this challenge with a brain-machine interface (BMI) ^29–32^ in which the transformation from neural activity to command was known exactly and determined by the experimenter. We trained rhesus monkeys to use motor cortex population activity to move a two-dimensional computer cursor on a screen through a BMI. The BMI transformed neural activity into a force-like command to update the cursor’s velocity, analogous to muscular force on the skeleton. Thus, an individual movement was produced by a series of commands, where each command acted on the cursor at an instant in time.

We discovered that the same exact command is issued with different neural activity patterns in different movements. Critically, these different patterns transition according to low-dimensional, invariant dynamics to patterns that issue the next command, even when the next command differs across movements. Thus, our results demonstrate that invariant dynamics drive commands to control different movements.

While past work has presented a view of how dynamics operate in a feedforward manner, propagating an initial state of activity ^23, 33, 34^ to produce movement, it has been unclear how feedback^24, 35–37^ integrates with invariant dynamics. Given that motor cortex is interconnected to larger motor control circuits including cortical^38–41^ and cortico-basal ganglia-thalamic circuits^8, 9, 42, 43^, we introduce a hierarchical model^44^ of optimal feedback control (OFC) in which the brain (i.e. larger motor control circuitry) uses feedback to control the motor cortex population which controls movement^45, 46^. Our model reveals that invariant dynamics can help transform feedback into commands, as they reduce the input that a population needs to issue commands. Altogether, our results demonstrate that invariant neural dynamics are both used and useful for issuing commands across different movements.

## Results

### BMI to study neural population control of movement

We used a BMI^47–49^ to study the dynamics of population activity as it issued commands for movement of a two-dimensional computer cursor (Fig. 1A). Population activity (20-151 units) was recorded using chronically implanted microwire electrode arrays spanning bilateral dorsal premotor cortex and primary motor cortex. Each unit’s spiking rate at time 𝑡 (computed as the number of spikes in a temporal bin) was stacked into a vector of population activity 𝑥_𝑡_, and the BMI used a “decoder” given by matrix 𝐾 to linearly transform population activity into a two-dimensional command:

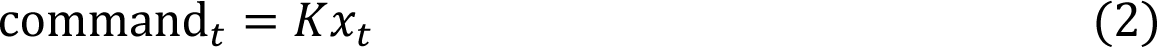

**Figure 1.**
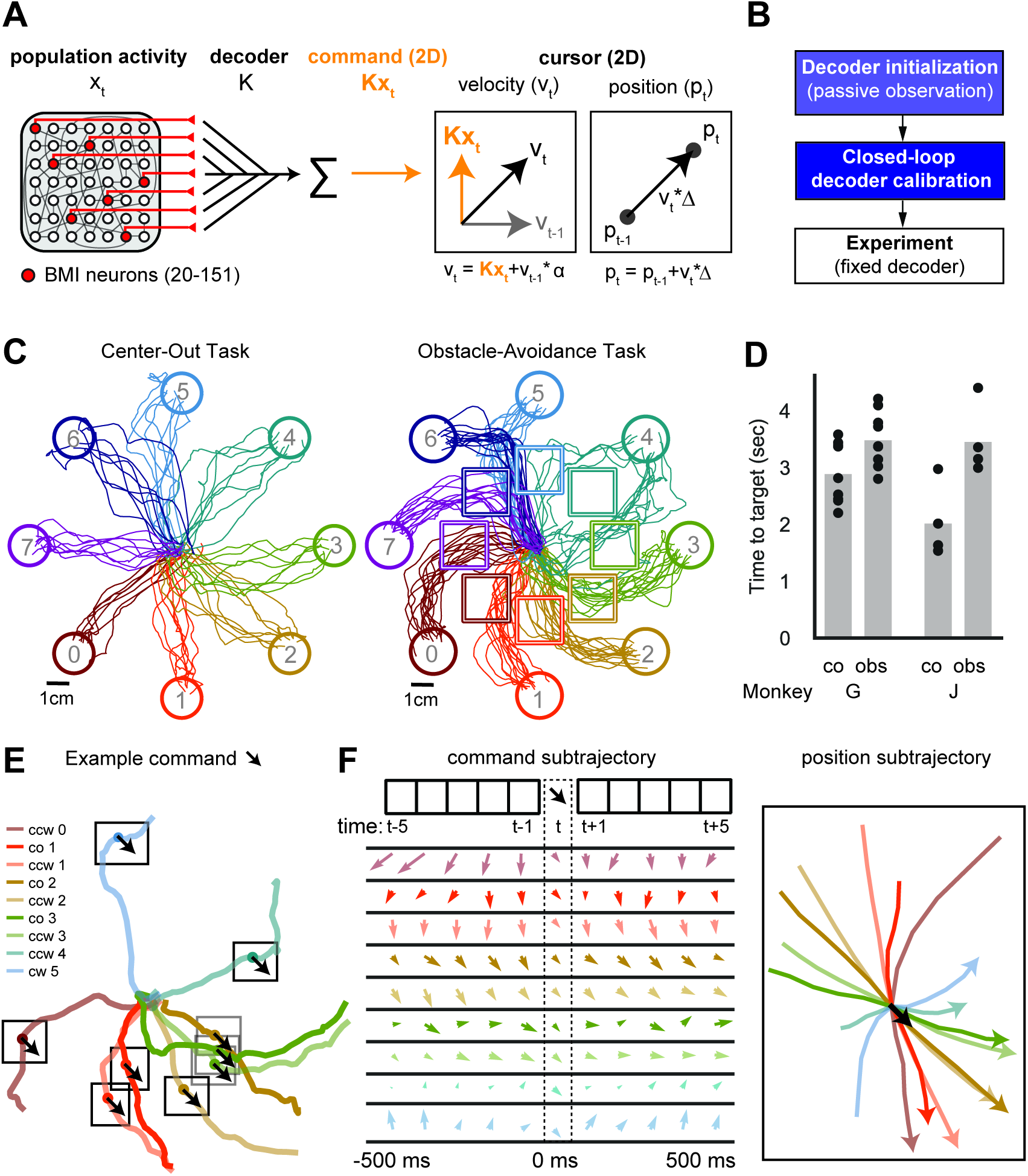
BMI to study neural population control of movement. (A) Schematic of the BMI system. (B) Schematic of decoder calibration. (C) Single trials of BMI control. (D) Average target acquisition time per session. (E) Example of the same command (black arrow) being issued during single trials of different conditions. The example command was in the -45 degree direction and the smallest magnitude bin of analysis. (F) *Left:* The average command subtrajectory from -500ms to 500ms. *Right:* The average position subtrajectory from -500ms to 500ms. See Fig. S1 for analysis of subtrajectories.

The command linearly updated the two-dimensional velocity vector of the computer cursor:

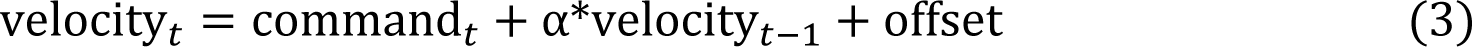

We note that the BMI was not identical across the two subjects, as neural activity was modeled with different statistical distributions (Gaussian for Monkey G and a Point Processs^47, 48^ for Monkey J, see STAR methods – “Neuroprosthetic decoding”).

The decoder was initialized as subjects passively watched cursor movement, calibrated as subjects used the BMI in closed-loop^49^ without performing trained overt movement, and then fixed for the experiment (Fig. 1B). Critically, the decoder was not fit during trained overt movement, as was done previously^16^, so it did not demand neural dynamics associated with overt movement.

To study control of diverse movements, we trained monkeys to perform two different tasks (Fig. 1CD). Monkeys performed a center-out task in which they moved the cursor from the center of the workspace to one of eight radial targets, and they performed an obstacle-avoidance task in which they avoided an obstacle blocking the straight path to the target. Our tasks elicited up to 24 conditions of movement (with an average of 16-17 conditions per session), where each condition is defined as the task performed (“co” = center-out task, “cw” / “ccw” = clockwise/counterclockwise movement around the obstacle in the obstacle-avoidance task) and the target achieved (numbered 0 through 7).

Importantly, the BMI enabled us to identify when neural activity issued the same exact command in different conditions (Fig. 1EF, Fig. S1). We considered two-dimensional, continuous-valued commands as the same if they fell within the same discrete bin for analysis. We categorized commands into 32 bins (8 angular x 4 magnitude) based on percentiles of the continuous-valued distribution (Fig. S1A; see STAR methods - “<u>Command discretization for analysis</u>”). On each session, a command (of the 32 discretized bins) was analyzed if it was used in a condition 15 or more times (Fig. S1B), for more than one condition. Each individual command was used with regularity during multiple conditions (on average ∼7 conditions, Fig. S1B), within distinct local “subtrajectories” (Fig. 1F, Fig. S1, STAR methods – “<u>Cursor and command trajectory visualization”</u>).

### Using the BMI to test whether invariant dynamics are used to control different movements

The BMI enabled us to test whether the pattern of neural activity systematically influences the subsequent pattern and command. We can visualize an activity pattern 𝑥_𝑡_ as a point in high-dimensional activity space, where each neuron’s activity is one dimension, and visualize the transition between two patterns 𝑥_𝑡_ and 𝑥_𝑡+1_ as an arrow (Fig. 2A). Then, dynamics can be visualized as a flow field in activity space. This flow field is invariant because the predicted transition for a given neural activity pattern (i.e. its arrow) does not change, regardless of the current command or condition. Because there are more neurons than dimensions of the command, different activity patterns can issue the same command^24, 50^ (Fig. 2B), as is believed to be true in the natural motor system^23, 24, 50^. The BMI decoder defined the “decoder space” as the dimensions of neural activity that determine the command and the “decoder null space” as the orthogonal dimensions which have no consequence on the decoder. The BMI allowed us to observe the precise temporal order of commands (Fig. 2C) and test whether activity trajectories followed the flow of invariant dynamics to issue these commands for movements (Fig. 2D).

**Figure 2.**
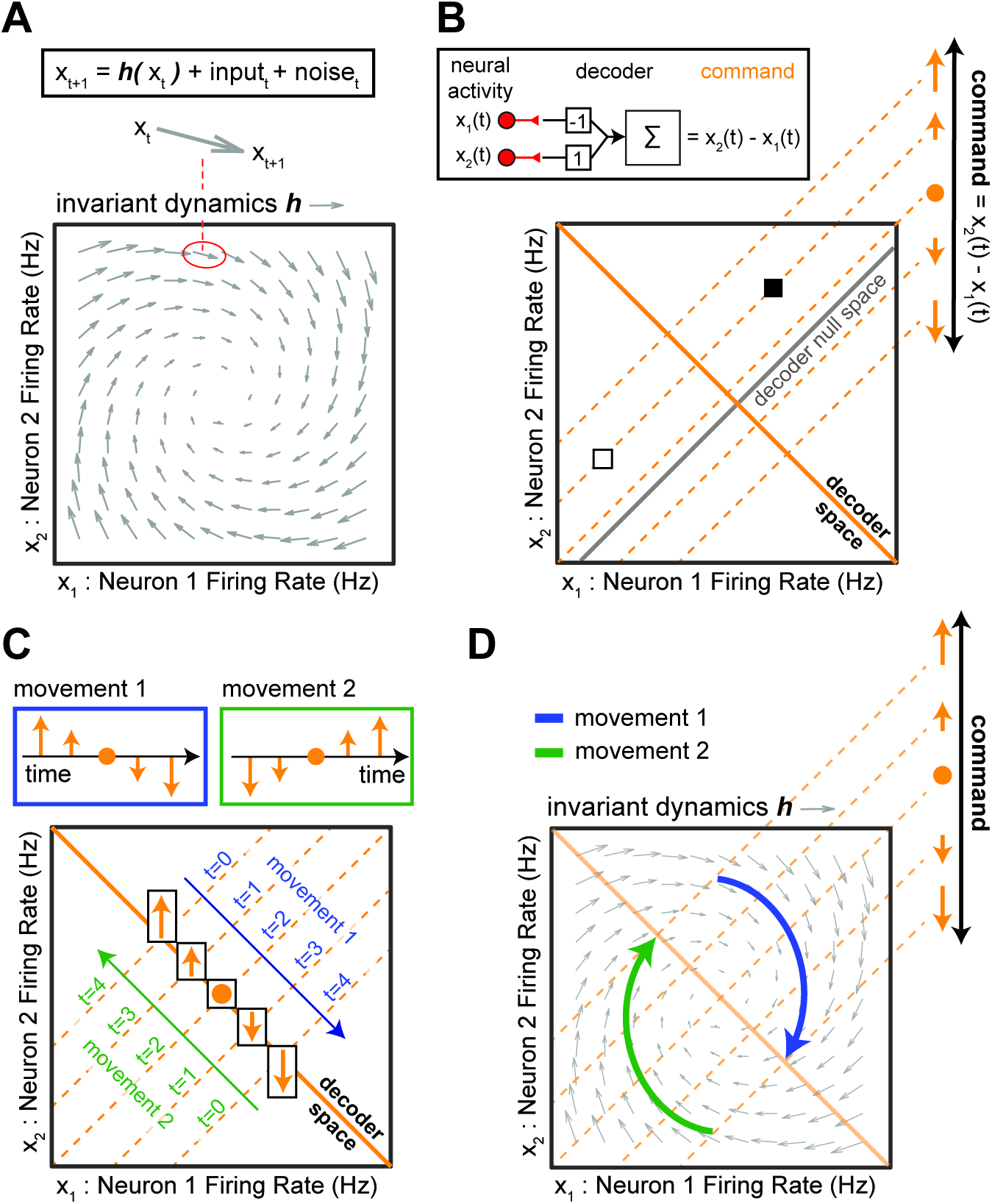
Using the BMI to test whether invariant dynamics are used to control different movements. (A) Illustration of invariant dynamics. (B) Multiple neural activity patterns (e.g. white and black square) issue the same command. An illustrative decoder defines the command at time 𝑡 as the difference between two neurons’ instantaneous activity 𝑥_2_(𝑡) − 𝑥_1_(𝑡), symbolized with orange arrows (top right) indicating the command’s magnitude and sign. (C) A trajectory of commands (orange arrows) produces one whole movement. Movement 1 (blue) and 2 (green) are driven by the same commands in different temporal orders. (D) Neural activity that follows invariant dynamics *h* in order to issue the commands for movement. See Fig. S3D for another example of invariant dynamics (decaying dynamics).

### The same command is issued by different neural activity patterns in different movements

First, we tested whether the same command is issued by different neural activity patterns in different movements, as would be expected if the current pattern influences the subsequent pattern and command (Fig. 3A). We calculated the distance between the average neural activity for a given command and condition and the average neural activity for the given command pooled over conditions. We then tested if this distance is larger than expected simply due to the variability of noisy neural activity. To emulate the scenario in which neural activity for a given command has the same distribution across conditions, we constructed shuffled datasets where we identified all observations of neural activity issuing a given command and shuffled their condition-labels, for all commands (see STAR methods – “<u>Behavior-preserving shuffle of activity</u>”). In this scenario, the distance is expected to be greater than zero simply because average activity is estimated from limited samples and thus is subject to variability.

**Figure 3.**
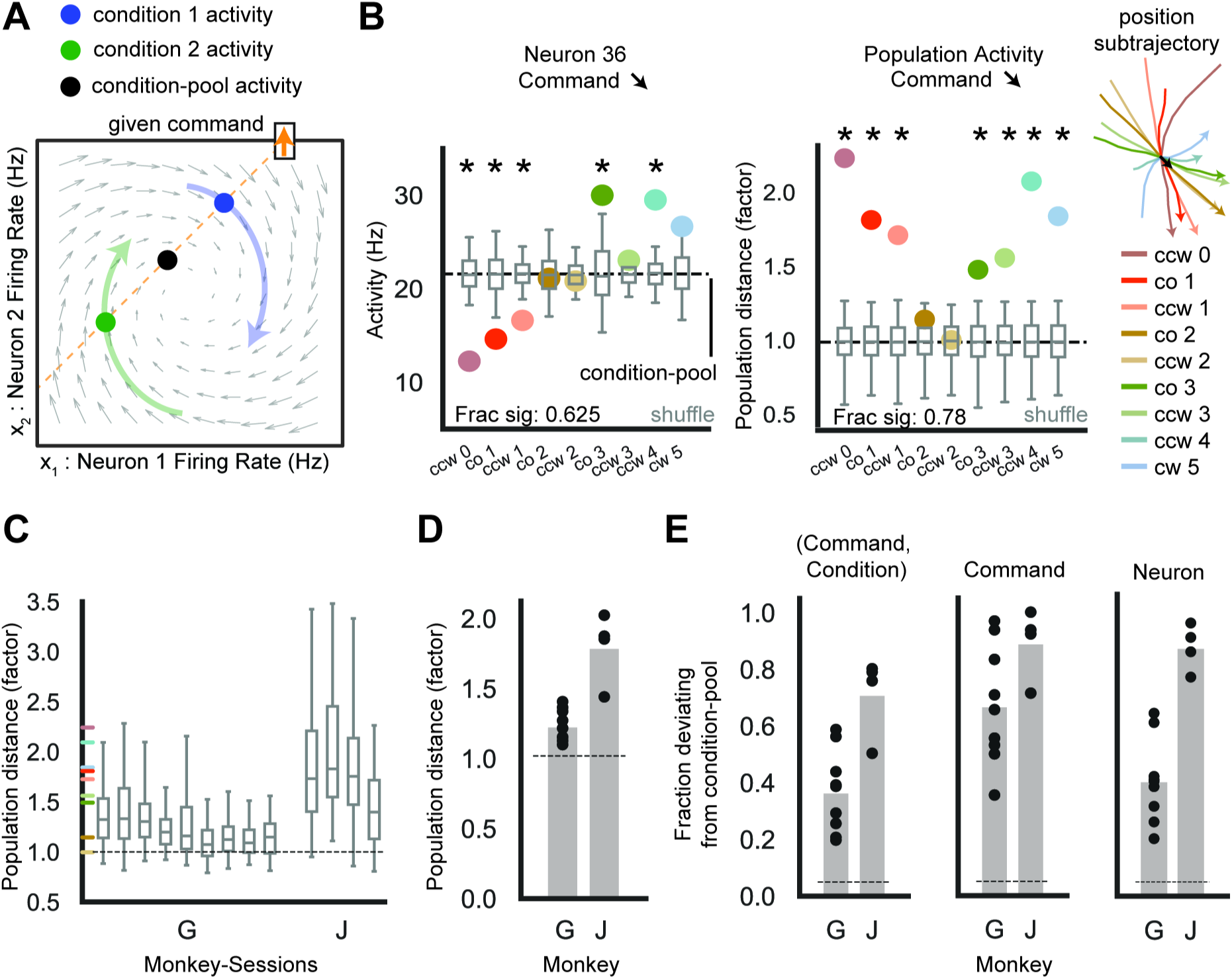
The same command is issued by different neural activity patterns in different movements. (A) The same command (orange upward arrow) is issued in different conditions with different activity patterns (blue, green dots). These patterns deviate from the condition-pooled average activity pattern for the command (black dot). (B) *Left:* An example neuron’s average firing rate (colored dots) for the example command and conditions from Fig. 1F (position subtrajectories plotted at right legend), as well as the condition-pooled average activity (dashed black line labeled “condition-pool”). The condition-shuffled distributions of average activity are shown with gray boxplots indicating the 2.5^th^, 25^th^, 50^th^, 75^th^, and 97.5^th^ percentiles. Asterisk indicates the distance for the (command, condition, neuron) exceeded the shuffle distance (p<0.05). 5/9 or 62.5% of the examples were significant. Distance was significantly greater than shuffle distance aggregating over all (command, condition, neuron) tuples: Monkey G [J]: p-value < 0.001 for 9/9 [4/4] sessions, p-value < 0.001 pooled over sessions. *Right:* Population distance normalized to the shuffle mean (colored dots). 7/9 or 78% of examples were significant. Fig. S2A shows population distances for all (command, condition) tuples in this session. (C) The distribution of normalized population distances across (command, condition) tuples. Colored ticks indicate distances in (B) *right*. See Fig. S2BC for additional distance distributions. (D) Normalized population distance averaged across (command, condition) tuples (Monkey G [J]: n=9 [4] sessions). Bars indicate the average across sessions. Population distance was significantly greater than shuffle distances, aggregating over all (command, condition) tuples: Monkey G [J]: p-value < 0.001 for 9/9 [4/4] sessions, p-value < 0.001 for pooled over sessions. (E) *Left:* Fraction of (command, condition) tuples with distance significantly greater than shuffle distance. *Middle:* Fraction of commands with distance significantly greater than shuffle distance, aggregating over conditions. *Right:* Fraction of neurons with distance significantly greater than shuffle distance, calculated for each (command, condition) separately and aggregating over all (command, condition) tuples for statistics. Throughout (E): dashed line indicates chance level (fraction equal to 0.05 significantly deviating from shuffle distance) and datapoints are each of 9 [4] sessions for monkey G [J]. See Fig. S6E-H for the relationship between population distance and command subtrajectories across pairs of conditions. See Table S1 for statistics details.

Overall, neural activity issuing a given command significantly deviated across conditions relative to the shuffle distribution (Fig. 3B-E). Distances averaged within-session ranged from 10% to 200% larger than shuffle distance (Fig. 3D, S2 for additional distributions). Distances were significantly larger than shuffle distances for a large fraction of individual (command, condition) tuples (∼30% for Monkey G, ∼70% for Monkey J), individual commands (∼65% for G, ∼90% for J) when aggregating over conditions, and individual neurons (∼40% for G, ∼80% for J) when aggregating over all (command, condition) tuples (Fig. 3E). Further, these deviations reflected the behavior; the distance between two patterns issuing the same command correlated with the distance between the command subtrajectories (Fig. S6E-H).

### Invariant dynamics predict the different neural activity patterns used to issue the same command

Given that a command was not issued with the same activity pattern across conditions, we next constructed a model of invariant dynamics. We used single-trial neural activity 𝑥_𝑡_ from all conditions to estimate dynamics with a linear model (Fig. 4A):

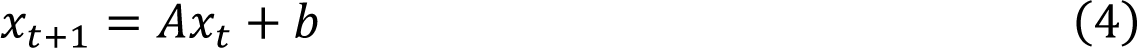

**Figure 4.**
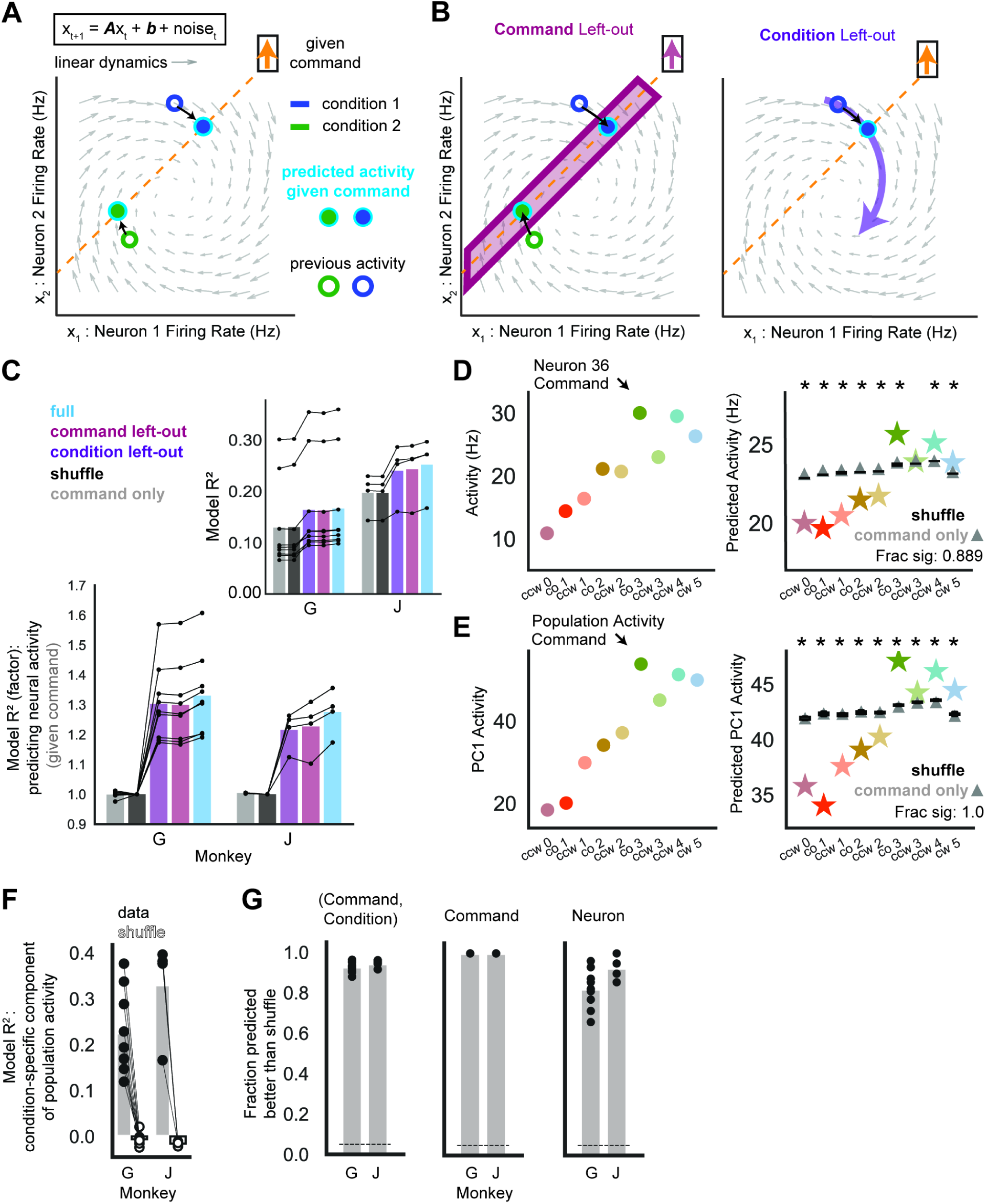
Invariant dynamics predict the different neural activity patterns used to issue the same command. (A) A linear dynamics model predicts the different activity patterns (cyan-outlined dots) that issue a given command (orange arrow) based on previous activity. See Fig. S6 for predictions of the relationship between activity patterns across pairs of conditions. (B) Models were tested on neural activity for a command (*Left*, magenta) or condition (*Right*, purple) left-out of training the model. See Fig. S4 for elaboration on invariant dynamics generalization. (C) The coefficient of determination (R^2^) of models predicting neural activity given the command it issues and previous activity, evaluated on test data not used for model fitting (Monkey G [J]: n=9 [4] sessions). See Fig. S3 for properties of the models. Inset shows raw R^2^, where “shuffle” is the 95^th^ percentile of the shuffle distribution of R^2^. Main panel shows R^2^ normalized to shuffle. Full dynamics, command left-out dynamics, and condition left-out dynamics all predicted neural activity significantly better than shuffle dynamics. For each model: Monkey G [J]: p-value < 0.001 for 9/9 [4/4] sessions, p-value < 0.001 for sessions pooled. Fig. S5 shows models with behavior variables and non-linear dynamics. (D) *Left.* Average activity for the example neuron, command, and conditions from Fig. 3B, left. *Right.* Prediction of the activity in *Left* by the full dynamics model (stars), the shuffle dynamics model (black boxplot distribution), and the model predicting neural activity only using the command (gray triangle). 8/9 or 88.9% of these examples were predicted significantly better than shuffle dynamics. The full dynamics model predicted individual neuron activity better than shuffle dynamics, aggregating over all (command, condition, neuron) tuples (Monkey G [J]: p-value < 0.001 for 9/9 [4/4] sessions, p-value < 0.001 for pooled sessions). (E) *Left.* Average population activity for the example command and conditions from Fig. 3B right, visualized along the activity dimension that captured the most variance (the first principal component, labeled “PC1”, of condition-specific average population activity). *Right.* Prediction of activity in *Left* by the full dynamics model (stars), the shuffle dynamics model (black boxplot distribution), and the model predicting neural activity only using the command (gray triangle). 9/9 or 100.0% of these examples were predicted with significantly lower error than shuffle dynamics (prediction was calculated using full population activity, not just PC1). The full dynamics model predicted population activity with lower error than shuffle dynamics, aggregating over all (command, condition, neuron) tuples (Monkey G [J]: p-value < 0.001 for 9/9 [4/4] sessions, p-value < 0.001 for pooled sessions). (F) Model R^2^ from predicting the component of average neural activity for a given command that is specific to a condition, comparing the full dynamics model (dark gray bar and filled dots) with the mean of the shuffle dynamics model (light bar and empty dots) (Monkey G [J]: n=9 [4] sessions). The full dynamics model predicted significantly more variance than shuffle dynamics (Monkey G [J]: p-value < 0.001 for 9/9 [4/4] sessions, p-value < 0.001 for pooled sessions). (G) *Left.* Fraction of (command, condition) tuples where full dynamics predicts average population activity significantly better than shuffle dynamics. *Center.* Fraction of commands where full dynamics predicts average population activity significantly better than shuffle dynamics, calculated for each condition separately and then aggregated over all conditions for statistics. *Right.* Fraction of neurons where full dynamics predicts the neuron’s average activity significantly better than shuffle dynamics, calculated for each (command, condition) separately and then aggregated over all (command, condition) tuples for statistics. Throughout E: datapoints are each of 9[4] sessions for Monkey G[J]. See Table S1 for statistics details.

We found that the dynamics 𝐴 were low-dimensional (∼4 dimensions, Fig. 5D, S3B) and decaying to a fixed point (Fig. S3A,C), contrasting with rotational dynamics observed during natural motor control ^12, 13, 16, 22, 51^. See Fig. S3D for an illustration of how decaying invariant dynamics can control different movements. Notably, a non-linear dynamics model (a recurrent switching linear dynamical system^52^) did not out-perform these simple linear dynamics (Fig. S5C-F).

**Figure 5.**
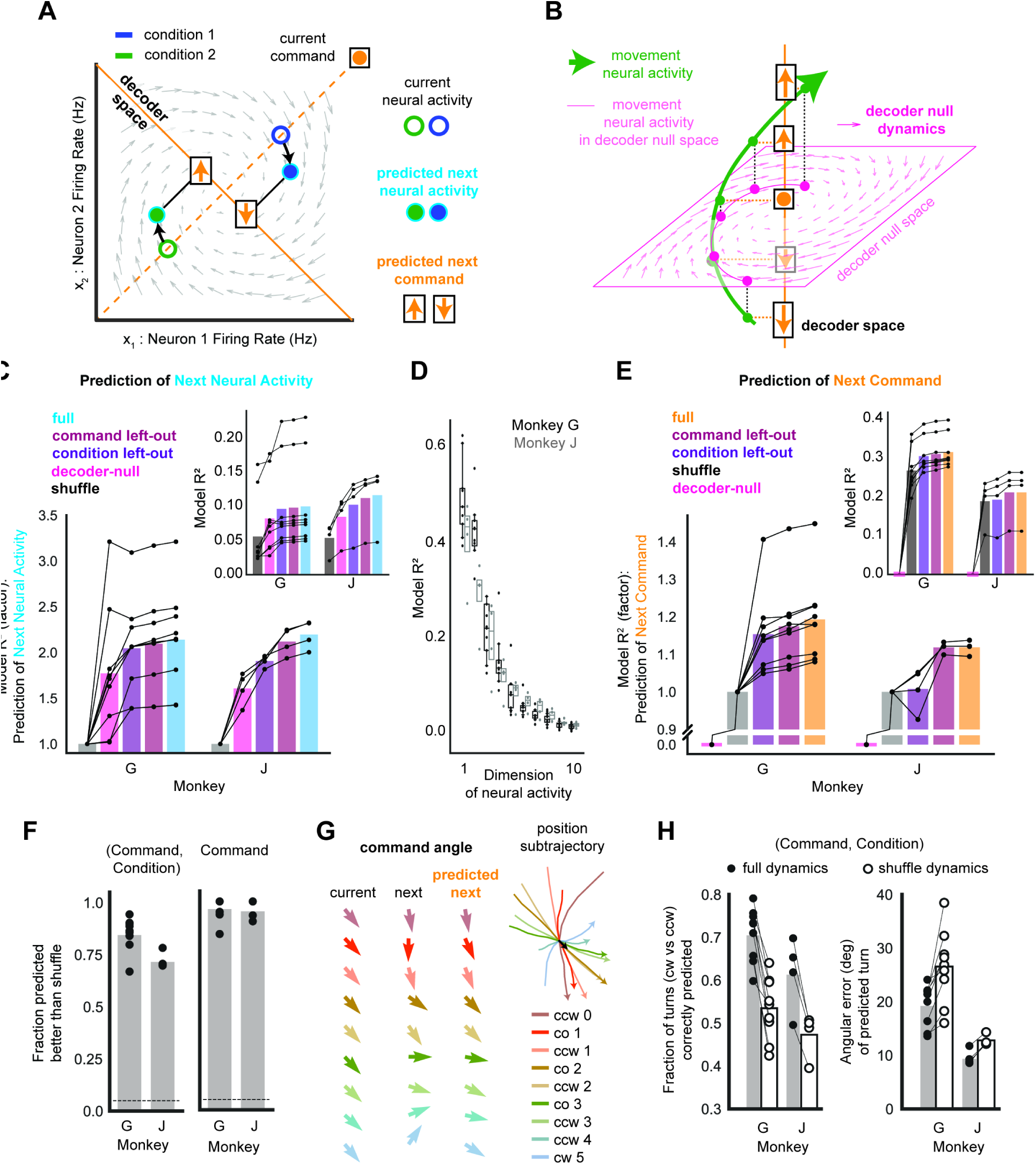
Invariant dynamics align with the decoder, propagating neural activity to issue the next command. (A) A linear dynamics model predicts the transition from current neural activity (colored rings) to next neural activity (cyan-outlined dots) and next commands (orange symbols) (i.e. the component of neural activity in the decoder space). (B) If invariant dynamics are low-dimensional and only occupy the decoder null space (pink plane), then they do not predict the next command (i.e. the component of neural activity in the decoder space). (C) The coefficient of determination (R^2^) of models predicting next neural activity given current neural activity, evaluated on test data not used for model fitting (Monkey G [J]: n=9 [4] sessions). Inset shows raw R^2^, where “shuffle” is the 95^th^ percentile of the shuffle distribution of R^2^. Main panel shows R^2^ normalized to shuffle. All models predicted next neural activity significantly better than shuffle dynamics. For each model, Monkey G [J]: p-value < 0.001 for 9/9 [4/4] sessions, p-value < 0.001 for sessions pooled. (D) R^2^ of full model for each neural activity dimension (dynamics eigenvector), sorted by R^2^. (E) Same as (C), except prediction of next command given current neural activity (Monkey G [J]: n=9 [4] sessions). All models except decoder-null dynamics predicted next command significantly better than shuffle dynamics. For condition left-out dynamics (purple), Monkey G[J]: p-value < 0.001 for 9/9 [2/4] session, p-value < 0.05 for 9/9 [3/4] session, p-value n.s. for 0/0 [1/4] sessions, p-value < 0.001 for sessions pooled. For full dynamics and command left-out dynamics, Monkey G [J]: p-value < 0.001 for 9/9 [4/4] sessions, p-value < 0.001 for sessions pooled. (F) Analyses of how well the next command is predicted for individual (command, condition) tuples. The full dynamics model predicted condition-specific next command better than shuffle dynamics, aggregating over all (command, condition) tuples (Monkey G [J]: p-value < 0.001 for 9/9 [4/4] sessions, p-value < 0.001 for pooled sessions). *Left.* Fraction of (command, condition) tuples where full dynamics predicts the next command significantly better than shuffle dynamics (Monkey G [J]: n=9 [4] sessions). *Right*. Fraction of commands where full dynamics predicts the next command significantly better than shuffle dynamics, calculated for each condition separately and then aggregated over all conditions for statistics (Monkey G [J]: n=9 [4] sessions). (G) Visualization of the command angle (*left*) (i.e. the direction that the command points) for the example command and conditions (*right*) from Fig. 3B. For each condition (each row), visualization shows the average current command angle (first column), the average next command angle (second column), and the prediction of the average next command angle by the full dynamics model (third column). (H) For each (command, condition) tuple, prediction of the angle between the next command and the condition-pooled average next command. *Left.* Fraction of (command, condition) tuples for which the sign of the angle is accurately predicted (positive=turn counterclockwise, negative=turn clockwise). Full dynamics predictions are significantly more accurate than shuffle dynamics (Monkey G [J]: p-value < 0.001 for 9/9 [4/4] sessions, p-value < 0.001 for pooled sessions. *Right.* Error in predicted angle. Full dynamics predictions are significantly more accurate than shuffle dynamics (Monkey G [J]: p-value < 0.001 for 9/9 [4/4] sessions, p-value < 0.001 for pooled sessions). See Table S1 for statistics details.

We asked whether invariant dynamics predict the different activity patterns observed to issue the same command. Concretely, we predicted the activity pattern given the command it issued and its previous activity (Fig. 4A, see STAR methods – “<u>Invariant dynamics model predictions</u>”), combining the dynamics model (Equation 4) with the decoder (Equation 2). This analyzed whether the model could predict the component of the activity pattern that can vary when a given command is issued, i.e. the component in the decoder null space. For comparison, we also computed the prediction of neural activity when only given the command it issued (in the absence of a dynamics model). Further, we tested whether the invariant dynamics model generalized to new commands and conditions. Dynamics models were fit on neural activity specifically excluding individual commands or conditions, and these models were used to predict the neural activity for the left-out commands or conditions (Fig. 4B, Fig. S4, see STAR methods – “<u>Invariant dynamics models</u>”).

We tested whether the dynamics model’s accuracy exceeded a dynamics model fit on the shuffled datasets that preserved the temporal order of commands while shuffling the neural activity issuing the commands (see STAR methods – “<u>Behavior-preserving shuffle of activity</u>”). The shuffle dynamics model captured the expected predictability in neural activity due to the predictability of commands in the performed movements.

On the level of single time points in individual trials, we found that the dynamics model significantly exceeded shuffle dynamics in predicting the activity pattern issuing a given command based on the previous pattern. Importantly, it generalized across left-out commands and conditions (Fig. 4C) and even when much larger subsets of commands and conditions were left-out (Fig. S4). We confirmed that the result was not driven by neural activity simply representing behavioral variables (cursor kinematics, target location, and condition) in addition to the command (Fig. S5AB), consistent with previous work ^53^.

The invariant dynamics model also predicted the different average activity patterns for each command and condition (Fig. 4D-G) significantly better than shuffle dynamics. It predicted 20-40% of the condition-specific component of neural activity (i.e. the difference between average activity for a (command, condition) and the prediction of that activity based on the command alone) (Fig. 4F, see STAR methods – “<u>Invariant dynamics model predictions</u>”). The model predicted neural activity for the vast majority of commands, conditions, and neurons (Fig. 4G), revealing that dynamics were indeed invariant.

Finally, the dynamics model preserved structure of neural activity across pairs of conditions (Fig. S6A-D) and predicted that the distance between two activity patterns issuing the same command would be correlated with the distance between the corresponding command subtrajectories (Fig. S6E-I). Altogether, these results show that invariant dynamics contribute to what activity pattern was used to issue a command, generalizing across commands and conditions.

### Invariant dynamics align with the decoder, propagating neural activity to issue the next command

We next asked whether invariant dynamics were actually used to transition between commands. Concretely, we used the dynamics model to predict the transition from the current activity pattern to the next pattern, and then we applied the BMI decoder to this prediction of next pattern in order to predict the next command (i.e. its continuous value) (Fig. 5A). This tests whether invariant dynamics predict the component of neural activity in the decoder space, which actually drives the BMI. The BMI enabled this analysis as it defines the transformation from neural activity to command which has not been measurable during natural motor control.

We emphasize that invariant dynamics do not have to predict the command, i.e. the decoder space (Fig. 5B). Low-dimensional dynamics could be misaligned with the decoder such that they only predict the component of neural activity in the decoder null space. To assess this possibility, we fit an invariant dynamics model on the component of neural activity in the decoder null space (“decoder-null dynamics”, see STAR methods – “<u>Invariant dynamics models</u>”). While this model was restricted to the decoder-null space, it maintained similar dimensionality and eigenvalues to the full dynamics model (Fig. S3BC).

Both the full dynamics and the decoder-null dynamics model predicted next neural activity significantly better than shuffle dynamics (Fig. 5C) on the level of single time points in individual trials. This reveals that invariant dynamics occupied decoder-null dimensions. Given that the full dynamics model was low-dimensional (Fig. S3B) and predicted ∼4 dimensions more accurately than the rest of neural activity (Fig. 5D), we next tested whether the dynamics aligned with the decoder. Critically, the full dynamics model predicted the next command (Fig. 5E) better than shuffle dynamics, while decoder-null dynamics provided absolutely no prediction for the next command, as expected by construction. The dynamics were invariant, as the full dynamics model generalized across commands and conditions that were left-out from model fitting (Fig. 5E) and predicted the next command for the majority of (command, condition) tuples (Fig. 5F). These predictions preserved structure across pairs of conditions, such that invariant dynamics indicated how similar the next command would be across pairs of conditions (Fig. S6I-K).

Notably, invariant dynamics could predict the turn that the next command would take following a given command in a specific condition relative to the average next command (averaged across conditions for the given current command) (Fig. 5GH). Specifically, the dynamics model predicted whether the turn would be clockwise or counter clockwise (Fig. 5H *left*) and the angle of turn (Fig 5H *right*) better than shuffle dynamics. Altogether, these results show that invariant dynamics align with the decoder and are used to transition between commands.

### An OFC model reveals that invariant dynamics reduce the input that a neural population needs to issue commands based on feedback

We observe that the invariant dynamics model did not perfectly predict transitions between commands. Throughout movement there were substantial residuals (Fig. S3E-G), consistent with ongoing movement feedback driving neural activity in addition to invariant dynamics. However, it has been unclear how the brain can integrate feedback with invariant dynamics to control movement. Thus, we constructed a model of optimal feedback control (OFC) that incorporates invariant neural dynamics.

We introduce a hierarchical model in which the brain controls the neural population which controls movement of the BMI cursor (Fig. 6A, Equation 5). Population activity 𝑥_𝑡_ issues commands for movement and is driven by three terms: invariant dynamics (which we hypothesize are intrinsic to some connectivity of the neural population), input, and noise. The brain transforms ongoing cursor state and population activity into the input to the population that is necessary to achieve successful movement. Concretely, the brain acts as an optimal linear feedback controller with knowledge of the neural population’s invariant dynamics, the BMI decoder, and the condition of movement. In this formulation, the brain’s objective was to achieve the target while using the smallest possible input to the population, which minimizes the communication from the brain to the population. Importantly, this incentivized the OFC model to optimize input in order to use invariant dynamics to control movement, rather than relying solely on input to issue commands. Consistent with this formulation, experiments show that thalamic input into motor cortex is optimized during motor learning^54^.

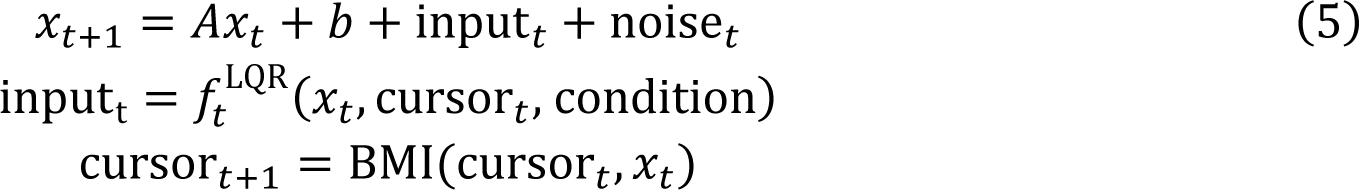

**Figure 6.**
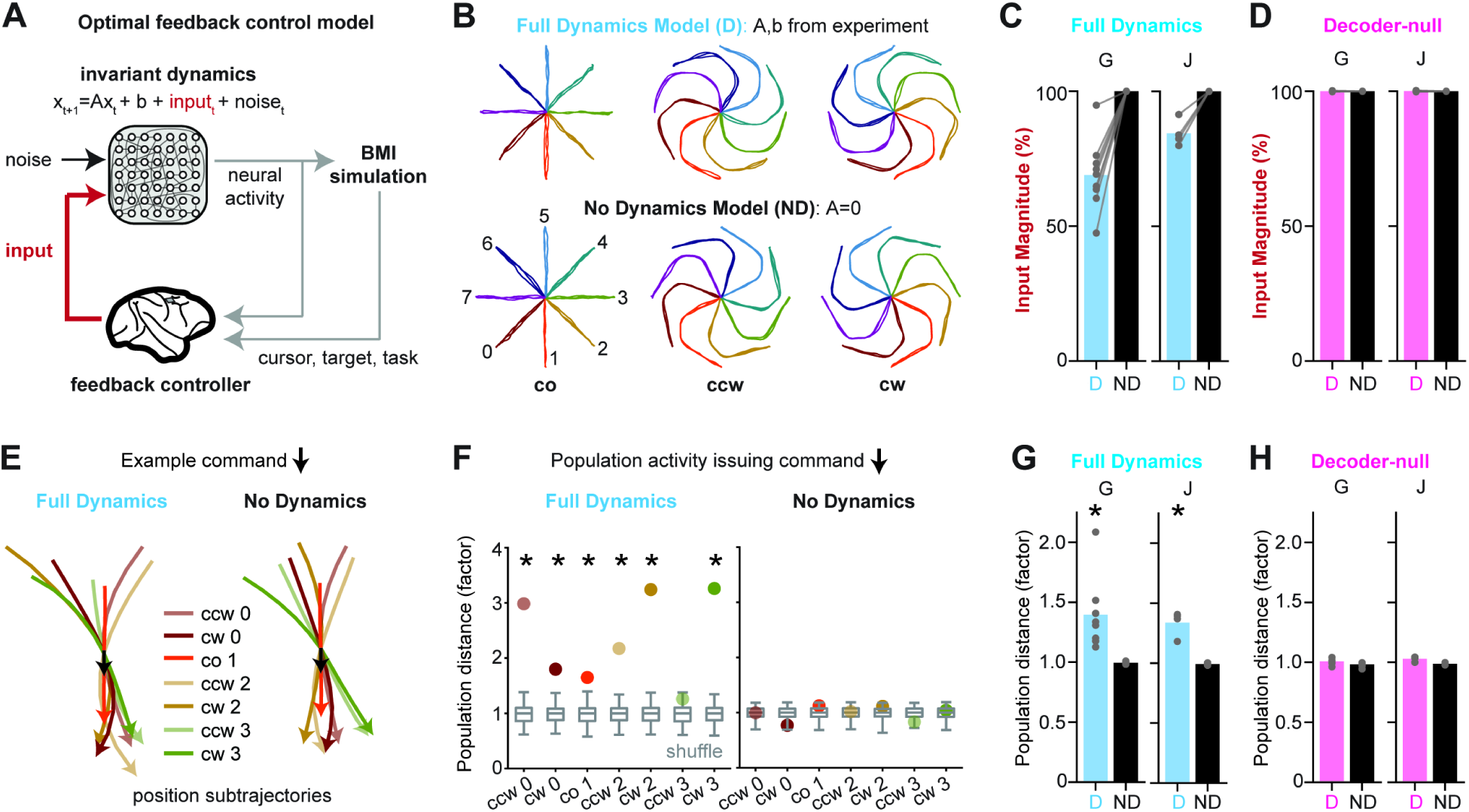
An OFC model reveals that invariant dynamics reduce the input that a neural population needs to issue commands based on feedback. (A) A model of optimal feedback control for movement that incorporates invariant neural dynamics. (B) Three simulated trials for each condition (center-out (co), counter-clockwise (ccw), and clockwise (cw) movements to 8 targets resulting in 24 conditions). *Top:* Full Dynamics Model that uses invariant dynamics fit on experimental data. *Bottom:* No Dynamics Model that uses dynamics matrix A set to 0. (C) Input magnitude as a percentage of the No Dynamics Model (Monkey G [J]: n=9 [4] sessions). The population required significantly less input to control movement under the Full Dynamics Model (cyan ‘D’) as compared to the No Dynamics Model (black ‘ND’). Un-normalized data were pooled across sessions and compared with a linear mixed effect (LME) model between input magnitude and model category with session modeled as random effect (Monkey G [J]: p-value < 0.001). Individual sessions were analyzed with a Wilcoxon signed-rank test that paired condition across the models (Monkey G [J]: p-value<0.05 for 9/9 [4/4] sessions). (D) Same as (C) but for Decoder-null Dynamics. There was no significant difference in input magnitude between Decoder-null Dynamics (pink ‘D’) and No Dynamics (black ‘ND’) when pooling across sessions (Monkey G [J] p-value > 0.05) and on individual sessions (Monkey G [J]: p-value<0.05 for 0/9 [0/4] sessions). (E) The same command is issued across conditions in both the Full Dynamics Model and No Dynamics Model. Average position subtrajectories are shown locked to an example command across conditions. (F) Distance between average population activity for a (command, condition) and the average activity for the command pooling across conditions, normalized by the mean distance of the shuffle distribution (gray boxplots showing mean, 0^th^ percentile, 25^th^, 75^th^, and 95^th^ percentile). *Left:* data from Full Dynamics Model. *Right:* data from the No Dynamics Model. Asterisk indicates distance is greater than shuffle (p-value<0.05). (G) Same as (F), but each point is an individual session pooling over (command, condition) tuples (Monkey G [J]: n=9 [4] sessions). Population distances for the Full Dynamics Model were greater than shuffle. Data was pooled over sessions using a LME with session modeled as random effect (Monkey G [J]: p-value < 0.001), and individual sessions were analyzed with a Mann-Whitney U test (p-value<0.05 for Monkey G [J] on 9/9 [4/4] sessions). No difference was detected in population distances between the No Dynamics Model and shuffle when pooling across sessions (Monkey G [J]: p-value > 0.05) and on individual sessions (p-value<0.05 for Monkey G (J) on 0/9 (0/4) sessions). (H) Same as (G), but for the Decoder-null Dynamics Model (pink ‘D’). No difference was detected in population distances between the Decoder-null Dynamics Model and shuffle when pooling across sessions (Monkey G [J]: p-value > 0.05) and on individual sessions (p-value<0.05 for Monkey G (J) on 0/9 (0/4) sessions). Also, no difference was detected in population distances between the No Dynamics Model and shuffle when pooling across sessions (Monkey G [J]: p-value > 0.05) and on individual sessions (p-value<0.05 for Monkey G(J) on 0/9 (0/4) sessions). See Table S2 for statistics details.

We simulated the model performing center-out and obstacle-avoidance movements with the decoders that were used in BMI experiments (see STAR methods – “<u>Optimal feedback control model and simulation</u>”). In the Full Dynamics Model, the brain computed the minimal input to a population that followed the invariant dynamics we observed experimentally. In the No Dynamics Model, the minimal input was computed to a neural population that had no invariant dynamics (i.e. the 𝐴 matrix was set to zero). To facilitate comparison, we designed the models to receive the same noise magnitude and to produce behavior with equal success and target acquisition time (Fig. 6B).

These simulations revealed that the population required significantly less input in the Full Dynamics Model than in the No Dynamics Model (Fig. 6C). This effect was erased in the Decoder-Null Dynamics Model (Fig 6D), in which the OFC model’s invariant dynamics were restricted to the decoder-null space. These results show that invariant dynamics that specifically align with the decoder, as experimentally-observed, can help the brain perform feedback control, reducing the input that the population needs to issue commands based on feedback.

Finally, we confirmed the principle that feedback control with invariant dynamics makes use of distinct activity patterns to issue a particular command. As in Fig. 3, we compared the OFC models’ neural activity against shuffled activity that preserved the temporal order of commands. The population activity distances for (command, condition) tuples were significantly larger than shuffle in the Full Dynamics Model but not in the No Dynamics Model (Fig. 6FG). Further, this effect depended on alignment between invariant dynamics and the decoder, as we detected no difference between the Decoder-Null Dynamics Model and shuffle (Fig. 6H). Thus, the OFC model used different neural activity patterns to issue the same command only when the invariant dynamics were useful for feedback control.

## Discussion

Theoretical work shows that recurrent connectivity can give rise to neural population dynamics for motor control^1, 4, 5^ and endow the brain with the capacity to generate diverse physical movement^3^. Experimental work has found that population activity in the motor cortex follows similar and predictable dynamics across different movements^11, 12, 16^. But it has been untested whether dynamics that are invariant across movements are used to actually control movement, as the transformation from neural activity to motor command has been challenging to measure^26, 27^ and model^23–25^. Here, we use a BMI to perform that test.

We discovered that different neural activity patterns are used to issue the same command in different movements. The activity patterns issuing the same command vary systemically depending on the past pattern, and critically, they transition according to low-dimensional, invariant dynamics towards activity patterns that causally drive the subsequent command. Our results’ focus on the command provides a conceptual advance beyond previous work that characterized properties of dynamics during behavior ^12, 13, 15, 16^, revealing that invariant dynamics are actually used to control movement.

Further, it has been unclear how the brain could integrate invariant dynamics with feedback ^24, 35–37^ to control movement. We introduce a hierarchical model^44^ of optimal feedback control, in which the brain uses feedback to control a neural population that controls movement. Optimal control theory reveals that invariant dynamics that are aligned to the decoder can help the brain perform feedback control of movement, reducing the input that a population needs to issue the appropriate commands. The model verified that when invariant dynamics are used for feedback control, the same command is issued with different neural activity patterns across movements. Altogether, these findings form a basis for future studies on what connectivity and neural populations throughout the brain give rise to invariant dynamics, whether the brain sends inputs to a neural population to take advantage of invariant dynamics, and whether invariant dynamics actually drive muscles during physical movement.

These results provide strong evidence against one traditional view that the brain reuses the same neural population activity patterns to issue a particular command. This perspective is present in classic studies that describe neurons as representing movement parameters^55, 56^. It is still debated what movement parameters are updated by motor cortex neurons ^28, 57–59^, as population activity encodes movement position ^60–62^, distance ^63^, velocity ^61, 62^, speed ^64^, acceleration ^65^, and direction of movement ^64, 66–68^, as well as muscle-related parameters such as force/torque ^55, 68–70^, muscle synergies ^71, 72^, muscle activation ^73–75^, and even activation of motor units^27^. Regardless of how commands from motor cortex update physical movement, our findings using a BMI strongly suggest that the motor cortex does not use the same neural activity pattern to issue a specific motor command. Our findings instead support the recent proposal that neural activity in motor cortex avoids “tangling”^11^ while issuing commands.

We found that invariant dynamics do not perfectly determine the neural population’s next command. We propose that as the brain sends input to the neural population, it performs feedback control on the state of the neural population’s invariant dynamics in order to produce movement. This proposal expands the number of behaviors for which invariant dynamics are useful. This is because invariant dynamics do not need to define the precise neural trajectories^12, 34^ that produce movement; they only need to provide useful transitions of neural activity that inputs can harness to control movement. In our data, simple dynamics (decaying dynamics with different time constants) in a low-dimensional activity space (∼4 dimensions) were used to control many conditions of movement (∼20 conditions). We find that invariant dynamics constrain neural activity in dimensions which do not directly matter for issuing current commands^50^, so that inputs in these dimensions can produce future commands (Fig. 6C). This mechanism refutes a simplistic interpretation of the minimal intervention principle^76^ in which neural activity should only be controlled in the few dimensions which directly drive commands. This also accords with the finding that motor cortex responses to feedback are initially in the decoder null space before transitioning to neural activity that issues corrective commands ^24^.

There is almost surely a limitation to the behaviors that particular invariant dynamics are useful for. Motor cortex activity occupies orthogonal dimensions and shows a different influence on muscle activation during walking and trained forelimb movement ^26^, and follows different dynamics for reach and grasp movements ^77^. Notably, our finding of decaying dynamics for BMI control contrasts with rotational dynamics observed during natural arm movement ^12, 13, 16, 22^. We speculate this could be because controlling the BMI relied more on feedback control than a well-trained physical movement, because controlling the BMI did not require the temporal structure of commands needed to control muscles for movement^2^, and/or because controlling the BMI did not involve proprioceptive feedback of physical movement^35^. Recent theoretical work shows that cortico-basal ganglia-thalamic loops can switch between different cortical dynamics useful for different temporal patterns of commands ^46^.

The use of invariant dynamics to issue commands has implications for how the brain learns new behavior ^78, 79^, enabling the brain to leverage pre-existing dynamics for initial learning ^25, 80, 81^ and to develop new dynamics through gradual reinforcement ^82, 83^. This learning that modifies dynamics relies on plasticity in cortico-basal ganglia circuits ^83–85^ and permits the brain to reliably access a particular neural activity pattern for a given command and movement ^32^, even if the same neural activity pattern is not used to issue the same command across different movements.

Modeling invariant dynamics can inform the design of new neuroprosthetics that can generalize commands to new behaviors ^16^ and classify entire movement trajectories ^86^. We expect that as new behaviors are performed, distinct neural activity patterns will be used to issue the same command, but that invariant dynamics can predict and thus recognize these distinct neural patterns as signal for the BMI rather than noise. In addition, our results inform the design of rehabilitative therapies to restore dynamics following brain injury or stroke to recover movement ^87, 88^.

Overall, this study put the output of a neural population into focus, revealing how rules for neural dynamics are used to issue commands and produce different movements. This was achieved by studying the brain as it controlled the very neural activity we recorded. BMI ^78, 89–92^, especially combined with technical advances in measuring, modeling, and manipulating activity from defined populations, provides a powerful technique to test emerging hypotheses about how neural circuits generate activity to control behavior.

## Acknowledgements

We thank I. Rodrigues-Vaz, D. Peterka, the Theory Center at the Zuckerman Institute, and I. Papusha for helpful discussions, and the Costa and Carmena labs for their support.

## Funding

NIH NINDS Pathway to Independence Award 1K99NS128250-01 (VRA)

BRAIN Initiative National Institute of Mental Health postdoctoral fellowship 1F32MH118714-01 (VRA)

NIH Pathway to Independence Award 1K99NS124748-01 (PK)

BRAIN Initiative National Institute of Mental Health postdoctoral fellowship 1F32MH120891-01 (PK)

NINDS/NIH BRAIN Initiative U19 NS104649 (RMC)

Simons-Emory International Consortium on Motor Control #717104 (RMC)

NINDS/NIH R01 NS106094 (JMC)

## Author contributions

V.R.A., P.K., R.M.C., and J.M.C. conceived and designed this study. P.K., S.G., and A.L.O. performed the experiments. P.K. and V.R.A. analyzed the data. All authors contributed materials and analysis tools. V.R.A., P.K., R.M.C, and J.M.C. wrote the manuscript. All authors reviewed the manuscript.

## Figures and legends

## STAR Methods

### RESOURCE AVAILABILITY

#### Lead contact

Further information and requests for resources and reagents should be directed to and will be fulfilled by the lead contacts, Rui M. Costa and Jose M. Carmena.

#### Materials availability

This study did not generate new unique reagents

#### Data and code availability

- Monkey BMI data (binned spike counts, cursor trajectories, condition parameters, decoder parameters, and task parameters) has been deposited the DANDI Archive at http://dandiarchive.org/dandiset/000404/draft and is publicly available as of the date of publication. Accession numbers / DOIs are listed in the key resources table.
- All original code has been deposited at https://github.com/pkhanna104/bmi_dynamics_code and is publicly available as of the date of publication. DOIs are listed in the key resources table.
- Any additional information required to reanalyze the data reported in this paper is available from the lead contact upon request.

### EXPERIMENTAL MODEL AND SUBJECT DETAILS

All training, surgery, and experimental procedures were conducted in accordance with the NIH Guide for the Care and Use of Laboratory Animals and were approved by the University of California, Berkeley Institutional Animal Care and Use Committee (IACUC). Two adult male rhesus macaque monkeys (7 years old, monkey G and 10 years old, monkey J) (Macaca mulatta, RRID: NCBITaxon:9544) were used as subjects in this study. Prior to this study, Monkeys G and J were trained at arm reaching tasks and spike-based 2D neuroprosthetic cursor tasks for 1.5 years. All animals were housed in pairs.

### METHOD DETAILS

#### Electrophysiology and experimental setup

Two male rhesus macaques were bilaterally, chronically implanted with 16 x 8 arrays of Teflon-coated tungsten microwire electrodes (35 mm in diameter, 500 mm separation between microwires, 6.5 mm length, Innovative Neurophysiology, Durham, NC) in the upper arm area of primary motor cortex (M1) and posterior dorsal premotor cortex (PMd). Localization of target areas was performed using stereotactic coordinates from a neuroanatomical atlas of the rhesus brain ^93^. Implant depth was chosen to target layer 5 pyramidal tract neurons and was typically 2.5-3 mm, guided by stereotactic coordinates.

During behavioral sessions, neural activity was recorded, filtered, and thresholded using the 128-channel Multichannel Acquisition Processor (Plexon, Inc., Dallas, TX) (Monkey J) or the 256-channel Omniplex D Neural Acquisition System (Plexon, Inc.) (Monkey G). Channel thresholds were manually set at the beginning of each session based on 1–2 min of neural activity recorded as the animal sat quietly (i.e. not performing a behavioral task). Single-unit and multi-unit activity were sorted online after setting channel thresholds. Decoder units were manually selected based on a combination of waveform amplitude, variance, and stability over time.

#### Neuroprosthetic decoding

Subjects’ neural activity controlled a two-dimensional (2D) neuroprosthetic cursor in real-time to perform center-out and obstacle-avoidance tasks. The neuroprosthetic decoder consists of two models:

1. A cursor dynamics model capturing the physics of the cursor’s position and velocity.
2. A neural observation model capturing the statistical relationship between neural activity and the cursor.

The neuroprosthetic decoder combines the models optimally to estimate the subjects’ intent for the cursor and to correspondingly update the cursor.

#### Decoder algorithm and calibration -- Monkey G

Monkey G used a velocity Kalman filter (KF) ^94, 95^ that uses the following models for cursor state 𝑐_𝑡_ and observed neural activity 𝑥_𝑡_ :

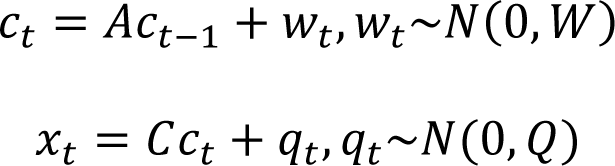

In the cursor dynamics model, the cursor state 𝑐_𝑡_ ∈ 𝑅^5^ was a 5-by-1 vector [*pos_x_*, *pox_y_*,*vel_x_vel_y_*, 1]^*T*^, 𝐴 ∈ 𝑅^5×5^ captures the physics of cursor position and velocity, and 𝑤_𝑡_ is additive Gaussian noise with covariance 𝑊 ∈ 𝑅^5𝑥5^ capturing cursor state variance that is not explained by 𝐴.

In the neural observation model, neural observation 𝑥_𝑡_ ∈ 𝑅^𝑁^ was a vector corresponding to spike counts from 𝑁 units binned at 10 Hz, or 100ms bins. 𝐶 models a linear relationship between the subjects’ neural activity and intended cursor state. The decoder only modeled the statistical relationship between neural activity and intended cursor velocity, so only the columns corresponding to cursor state velocity and the offset (columns 3-5) in 𝐶 were non-zero. 𝑄 is additive Gaussian noise capturing variation in neural activity that is not explained by 𝐶𝑐_𝑡_. For Monkey G, 35-151 units were used in the decoder (median 48 units).

In summary, the KF is parameterized by matrices {𝐴 ∈ 𝑅^5𝑥^^5^, 𝑊 ∈ 𝑅^5𝑥5^, 𝐶 ∈ 𝑅^𝑁𝑥5^, 𝑄 ∈ 𝑅^𝑁𝑥𝑁^}. The KF equations used to update the cursor based on observations of neural activity are defined as in ^95^.

The KF parameters were defined as follows. For the cursor dynamics model, the 𝐴 and 𝑊 matrices were fixed as in previous studies ^96^. Specifically, they were:

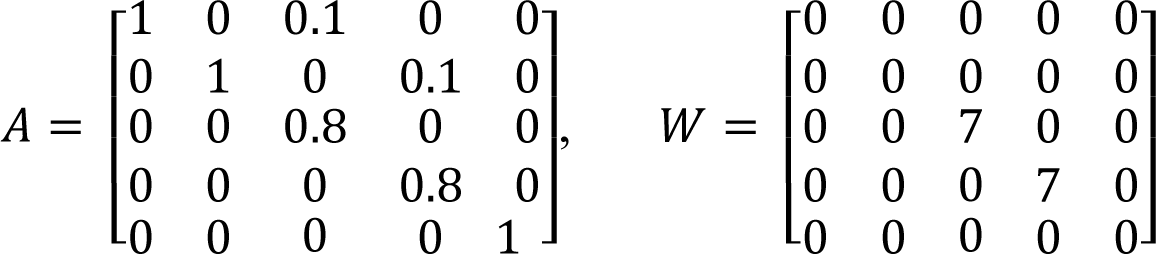

where units of cursor position were in cm and cursor velocity in cm/sec.

For the neural observation model, the 𝐶 and 𝑄 matrices were initialized from neural and cursor kinematic data collected at the beginning of each experimental session while Monkey G observed 2D cursor movements that moved through either a center-out task or obstacle avoidance task. Maximum likelihood methods were used to fit 𝐶 and 𝑄.

Next, Monkey G performed a “calibration block” where he performed the center-out or obstacle-avoidance task movements as the newly initialized decoder parameters were continuously calibrated/adapted online (“closed-loop decoder adaptation”, or CLDA). This calibration block was performed in order to arrive at parameters that would enable excellent neuroprosthetic performance. Every 100ms, decoder matrices 𝐶 and 𝑄 were adapted using the recursive maximum likelihood CLDA algorithm ^49^. Half-life values, defining how quickly 𝐶 and 𝑄 could adapt, were typically 300 sec, and adaptation blocks were performed with a weak, linearly decreasing “assist” (re-defining 𝑐_𝑡_ as a weighted linear combination of user-generated 𝑐_𝑡_ and optimal 𝑐_𝑡_ to drive the cursor to the target). Typical assist values at the start of the block were 90% user-generated, 10% optimal and decayed to 100% user-generated, 0% optimal over the course of the block. Following CLDA, decoder parameters were fixed. Then the experiment proceeded with Monkey G performing the center-out and obstacle-avoidance tasks.

#### Decoder algorithm -- Monkey J

Monkey J used a velocity Point Process Filter (PPF) ^47, 48^. The PPF uses the same cursor dynamics model for cursor state 𝑐_𝑡_ as the KF above, but uses a different neural observations model (a Point Process model rather than a Gaussian model) for the spiking *S*_*t*_^1:*N*^ of each of 𝑁 neurons:

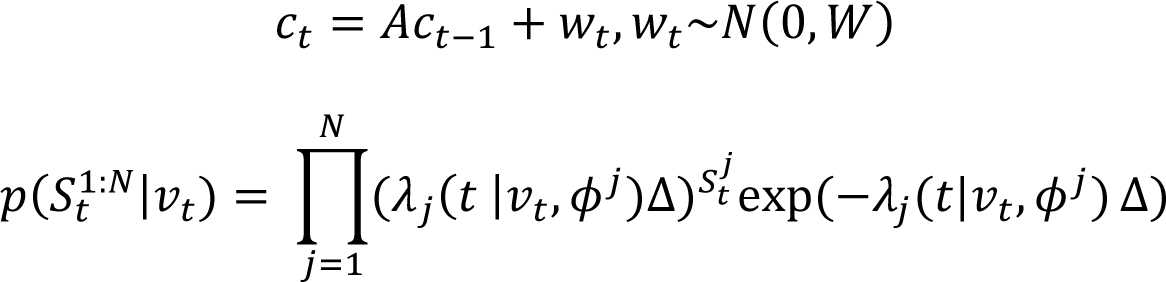

In the neural observations model, neural observation *S_t_^j^* is the j^th^ neuron’s spiking activity, equal to 1 or 0 depending on whether the j^th^ neuron spikes in the interval (𝑡, 𝑡 + Δ). We used Δ𝑡 = 5ms bins since consecutive spikes rarely occurred within 5ms of each other. For Monkey J, 20 or 21 units were used in the decoder (median 20 units). The probability distribution over spiking 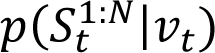 was a point process with 𝜆_𝑗_(𝑡 |𝑣_𝑡_, 𝜙^𝑗^) as the j^th^ neuron’s instantaneous firing rate at time t. 𝜆_𝑗_(𝑡 |𝑣_𝑡_, 𝜙^𝑗^) depended on the intended cursor velocity 𝑣_𝑡_ ∈ 𝑅^2^ in the two dimensional workspace and the parameters 𝜙^𝑗^ for how neuron *j* encodes velocity. 𝜆_𝑗_(𝑡 |𝑣_𝑡_, 𝜙^𝑗^) was modeled as a log-linear function of velocity:

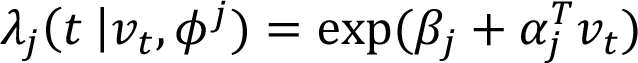

where 𝜙^𝑗^ parameters consist of 𝛼_𝑗_ ∈ 𝑅^2^, 𝛽_𝑗_ ∈ 𝑅^1^.

In summary, the PPF is parameterized by 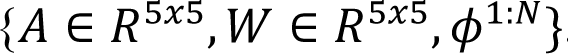. The PPF equations used to update the cursor based on observations of neural activity are defined as in ^48^.

The PPF parameters were defined as follows. For the cursor dynamics model, the 𝐴 and 𝑊 matrices are defined as:

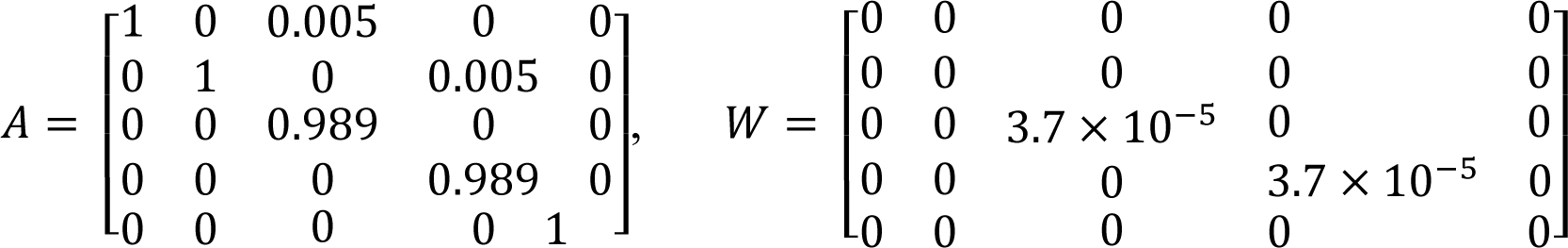

where units of cursor position were in m and cursor velocity in m/sec.

For the neural observations model, parameters 𝜙^1:𝑁^ were initialized from neural and cursor kinematic data collected at the beginning of each experimental session while Monkey J observed 2D cursor movements that moved through a center-out task. Decoder parameters were adapted using CLDA and optimal feedback control intention estimation as outlined in ^47^. Following CLDA, decoder parameters were fixed. Then the experiment proceeded with Monkey J performing the center-out and obstacle-avoidance tasks.

#### Definition of the command for the BMI

We defined the “command” for the BMI as the direct influence of subjects’ neural activity 𝑥_𝑡_ (binned at 100ms) on the cursor. Concretely, in both decoders, the command was a linear transformation of neural activity that we write as 𝐾𝑥_𝑡_ which updated the cursor velocity.

##### Command definition -- Monkey G

For Monkey G, the update to the cursor state 𝑐_𝑡_ due to cursor dynamics and neural observation 𝑥_𝑡_ can be written as:

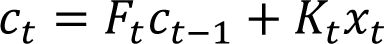

where 𝐹_𝑡_𝑐_𝑡−1_ is the update in cursor state due to the cursor dynamics process and 𝐾_𝑡_𝑥_𝑡_ is what we have defined as the command: the update in cursor state due to the current neural observation. 𝐾_𝑡_ ∈ 𝑅^5𝑥𝑥^ is the Kalman Gain matrix and 𝐹_𝑡_ = (𝐼𝐼 − 𝐾_𝑡_𝐶)𝐴. In practice 𝐾_𝑡_ converges to its steady-state form 𝐾 within a matter of seconds ^97^, and thus 𝐹_𝑡_ converges to 𝐹 = (𝐼𝐼 − 𝐾𝐶)𝐴, so we can write the above expression in its steady state form:

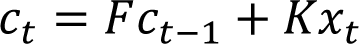

In our implementation, the structure of 𝐾 is such that neural activity 𝑥_𝑡_ directly updates cursor velocity, and velocity integrates to update position. The following technical note explains the structure of 𝐾. Due to the form of the 𝐴, 𝑊 matrices, *Rank*(𝐾) = 2. In addition, decoder adaptation imposed the constraint that the intermediate matrix 𝐶^𝑇^𝑄^−1^𝐶 was of the form a𝐼, where *a* = *mean*(*diag*(𝐶^𝑇^𝑄^−1^𝐶)). Due to this constraint, the rows of 𝐾 that update the position of the cursor are equal to the rows of 𝐾 that update the velocity multiplied by the update timestep: 𝐾(1: 2, ∶) = 𝐾(3: 4, ∶) ∗ 𝑑𝑡 ^98^ (see independent velocity control in the reference). Given this structure of 𝐾, neural activity’s contribution to cursor position is the simple integration of neural activity’s contribution to velocity over one timestep.

In summary, since 𝐾𝑥_𝑡_ reflects the direct effect of the motor cortex units on the velocity of the cursor, we term the velocity components of 𝐾𝑥_𝑡_ the “command”. We analyzed the neural spike counts binned at 100ms that were used online to drive cursor movements with no additional pre-processing.

##### Command definition -- Monkey J

For Monkey J the cursor state updates in time as:

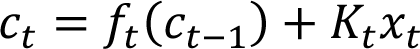

where

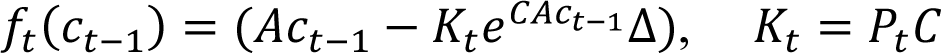

Here 𝑓_𝑡_(𝑐_𝑡−1_) is the cursor dynamics process and 𝐾_𝑡_𝑥_𝑡_ is the neural command. 𝑃_𝑡_ ∈ 𝑅^5𝑥5^ is the estimate of cursor state covariance, and 𝐶 ∈ 𝑅^5𝑥𝑁^ captures how neural activity encodes velocity as a matrix where each column is composed of 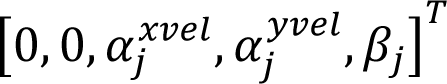 for the *j*th unit.

We define the command for analysis in this study as 𝐾_𝑥𝑒𝑡_𝑥_𝑡_, where 𝐾_𝑥𝑒𝑡_ is a time-invariant matrix that almost perfectly approximates 𝐾_𝑡_. While the PPF’s 𝐾_𝑡_ does not necessarily converge in the same way it does in the KF, for all four analyzed sessions, neural activity mapped through 𝐾_𝑥𝑒𝑡_ ∈ 𝑅^2𝑥𝑁^ could account for 99.6, 99.6, 99.5, and 99.8 percent of the variance of the command respectively (𝐾_𝑡_𝑥_𝑡_ ≅ 𝐾_𝑥𝑒𝑡_𝑥_𝑡_). In addition, due to the accuracy of this linear approximation, we also match Monkey J’s timescale of neural activity and commands to that of Monkey G. In order to match timescales across the two animals (Monkey G: 100 ms updates, Monkey J: 5ms updates), Monkey J’s commands were aggregated into 100 ms bins by summing 𝐾_𝑥𝑒𝑡_𝑥_𝑡_ over 20 consecutive 5ms bins to yield the aggregated command over 100ms. Correspondingly, Monkey J’s neural activity was also summed into 100ms bins by summing 𝑥_𝑡_ over 20 consecutive 5ms bins.

#### Neuroprosthetic tasks

Subjects performed movements in a two-dimensional workspace (Monkey J: 24cm x 24cm, Monkey G: 50cm x 28cm) for two neuroprosthetic tasks: a center-out task and an obstacle-avoidance task. We define the movement “condition” as the task performed (“co” = center-out task, “cw” / “ccw” = clockwise/counterclockwise movement around the obstacle in the obstacle-avoidance task) and the target achieved (numbered 0 through 7). Thus, there were up to 24 different conditions possible (8 center-out conditions, 8 clockwise obstacle-avoidance conditions, 8 counterclockwise obstacle-avoidance conditions). In practice, subjects mostly circumvented the obstacles for a given target location consistently in a clockwise or counterclockwise manner (as illustrated in Fig. 1C right) resulting in an average of 16-17 conditions per session.

##### Center-out task

The center-out task required subjects to hold their cursor within a center target (Monkey J: radius = 1.2 cm, Monkey G: radius = 1.7 cm) for a specified period of time (Monkey J: hold = 0.25 sec, Monkey G: hold = 0.2 sec) before a go cue signaled the subjects to move their cursor to one of eight peripheral targets uniformly spaced around a circle. Each target was equidistant from the center starting target (Monkey J: distance = 6.5cm, Monkey G: distance = 10cm). Subjects then had to position their cursor within the peripheral target (Monkey J: target radius = 1.2cm, Monkey G: target radius = 1.7cm) for a specified period to time (Monkey J: hold = 0.25, Monkey G: hold = 0.2sec). Failure to acquire the target within a specified window (Monkey J: 3-10 sec, Monkey G: 10 sec) or to hold the cursor within the target for the duration of the hold period resulted in an error. Following successful completion of a target, a juice reward was delivered. Monkey J was required to move his cursor back to the center target to initiate a new trial, and Monkey G’s cursor was automatically reset to the center target to initiate a new trial.

##### Obstacle-avoidance task

Monkey G performed an obstacle-avoidance task with a very similar structure to the center-out task. The only difference was that a square obstacle (side length 2 or 3 cm) would appear in the workspace centered exactly in the middle of the straight line connecting the center target position and peripheral target position. If the cursor entered the obstacle, the trial would end in an error, and the trial was repeated.

Monkey J’s obstacle-avoidance task required a point-to-point movement between an initial (not necessarily center) target and another target. On arrival at the initial target, an ellipsoid obstacle appeared on the screen. If the cursor entered the obstacle at any time during the movement to the peripheral target, an error resulted, and the trial was repeated. Target positions and obstacle sizes and positions were selected to vary the amount of obstruction, radius of curvature around the obstacles, and spatial locations of targets. Trials were constructed to include the following conditions: no obstruction, partial obstruction with low-curvature, full obstruction with a long distance between targets, and full obstruction with a short distance between targets thus requiring a high curvature. See ^48^ for further details. In this study, only trials that included partial obstruction or full obstruction were analyzed as “obstacle-avoidance” trials.

##### Number of sessions

We analyzed 9 sessions of data from Monkey G and 4 sessions of data from Monkey J where on each session, monkeys performed both the center-out and obstacle-avoidance tasks with the same decoder. Only successful trials were analyzed.

#### Optimal feedback control model and simulation

We introduce a model based on optimal feedback control (OFC) for how the brain can use invariant neural population dynamics to control movement based on feedback. From the perspective of the brain trying to control the BMI, we used the model to ask how invariant neural population dynamics affect the brain’s control of movement.

Thus, we performed and analyzed simulations of a model in which the brain acts as an optimal linear feedback controller (finite horizon linear quadratic regulator), sending inputs to a neural population so that it performs the center-out and obstacle-avoidance tasks (Fig. 6). The feedback controller computed optimal inputs to the neural population based on the current cursor state and current neural population activity. Specifically, the inputs were computed as the solution of an optimization problem that used knowledge of the target and task, decoder, and the neural population’s invariant dynamics. We simulated 20 trials for each of 24 conditions: 8 center-out conditions, 8 clockwise obstacle-avoidance conditions, and 8 counterclockwise obstacle-avoidance conditions. The neural and cursor dynamics processes in the simulation are summarized below:

##### Neural population dynamics with input

In our simulation, the neural activity of 𝑁 neurons 𝑥_𝑡_ ∈ 𝑅^𝑁^ is driven by invariant dynamics 𝐴 ∈ 𝑅^𝑁×𝑁^ that act on previous activity 𝑥_𝑡−1_, an activity offset 𝑏 ∈ 𝑅^𝑁^, inputs from the feedback controller 𝑢_𝑡−1_ ∈ 𝑅^𝑁^ that are transformed by input matrix 𝐵 ∈ 𝑅^𝑁×𝑁^, and noise 𝜎_𝑡−1_ ∈ 𝑅^𝑁^:

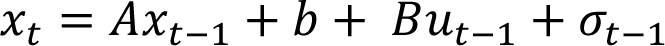

The input matrix 𝐵 was set to be the identity matrix such that each neuron has its own independent input. Each neuron also had its own independent, time-invariant noise (see *Noise* section below for how the noise level was set).

For notational convenience, an offset term was appended to 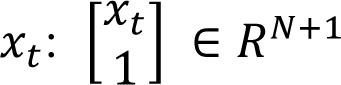. This enabled incorporating the offset 𝑏 into the neural dynamics matrix:

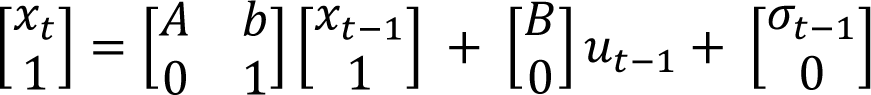

##### BMI cursor dynamics

The cursor update equations for the simulation matched the steady state cursor update equations in the online BMI experiment (see “<u>Definition of the command for the BMI</u>” above):

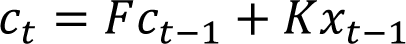

As in the experiment, cursor state 𝑐_𝑡_ ∈ 𝑅^𝑁𝑐^ where 𝑁_𝑐_ = 5 was a vector consisting of two-dimensional position, velocity, and an offset: [*pos_x_*, *pox_y_*, *vel_x_*, *vel_y_*, 1]^*T*^. *K* ϵ *R^N_c_×N^* was the decoder’s steady-state Kalman gain (Monkey G) or estimated equivalent 𝐾_𝑥𝑒𝑡_ (Monkey J). 𝐹 ∈ 𝑅^𝑁𝑐×𝑁𝑐^ was set to the decoder’s steady-state cursor dynamics matrix (Monkey G). For Monkey J, 𝐹 was estimated using the expression for calculating the steady-state cursor dynamics matrix: 𝐹_𝑥𝑒𝑡_ = (𝐼𝐼 − 𝐾_𝑥𝑒𝑡_𝐶_𝑥𝑒𝑡_) ∗ 𝐴_100𝑚𝑚𝑒_, where 𝐼𝐼 ∈ 𝑅^𝑁𝑐𝑥𝑁𝑐^, 𝐶_𝑥𝑒𝑡_ ∈ 𝑅^𝑁𝑥𝑁𝑐^ was set using the 𝛼, 𝛽 velocity encoding parameters from the point process filter (see above): 𝐶_𝑥𝑒𝑡_(𝑗, ∶) = [0 0 0.01 ∗ 𝛼_𝑗_(1) 0.01 ∗ 𝛼_𝑗_(2) 0.01 ∗ 𝛽_𝑗_]. Values in 𝐶_𝑥𝑒𝑡_ were multiplied by 0.01 to adjust for velocities expressed in units of cm/sec (in the simulation) instead of m/sec (as in PPF). 𝐴_100𝑚*s*_ was set to the same 𝐴 used by Monkey G so that the cursor dynamics would be appropriate for 100ms timesteps:

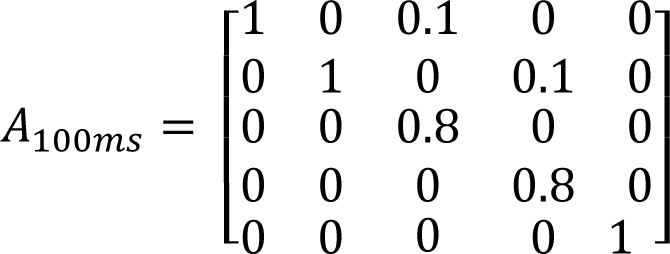

##### Joint dynamics of neural activity and cursor

The feedback controller sent inputs to the neural population which were optimal considering the task goal, the cursor’s current state, the neural population’s invariant dynamics, and the neural population’s current activity. To solve for the optimal input given all the listed quantities, first, the neural and cursor states are jointly defined. We append the cursor state 𝑐_𝑡_ to the neural activity state 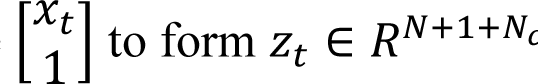:

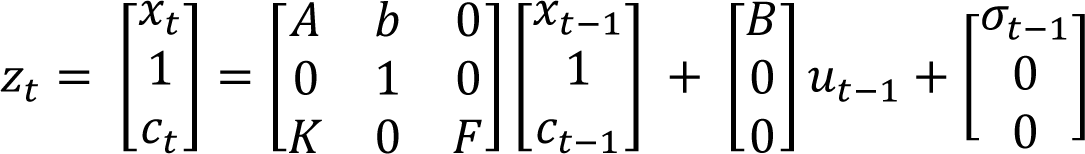

In words, this expression defines a linear dynamical system where input 𝑢_𝑡−1_ influences only the neural activity 𝑥_𝑡_, 𝑥_𝑡_ evolves by invariant dynamics 𝐴 with offset vector 𝑏, and 𝑥_𝑡_ drives cursor 𝑐_𝑡_ through the BMI decoder 𝐾. Finally, noise 𝜎_𝑡−1_ only influences neural activity 𝑥_𝑡_ (see *Noise* section below for how the noise level was set).

##### OFC to reach a target

Our OFC model computes input 𝑢_𝑡_ to the neural population such that the activity of the neural population 𝑥_𝑡_ drives the cursor to achieve the desired final cursor state (i.e. the target) with minimal magnitude of input 𝑢_𝑡_. Concretely, in the finite horizon LQR model, the optimal control sequence (𝑢_𝑡_, 𝑡 = 0, 1, … 𝑇 − 1) is computed by minimizing the following cost function:

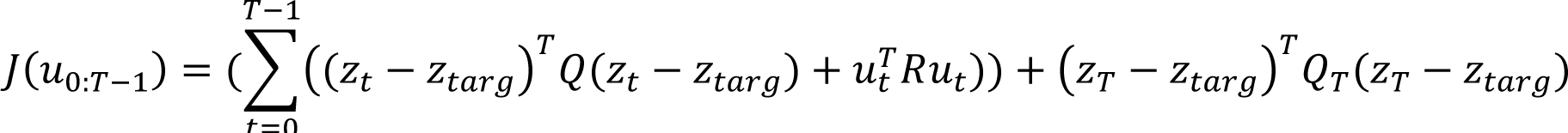

In our model, 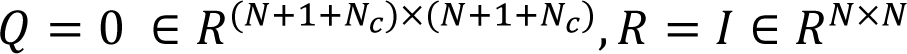, and *Q_T_* = 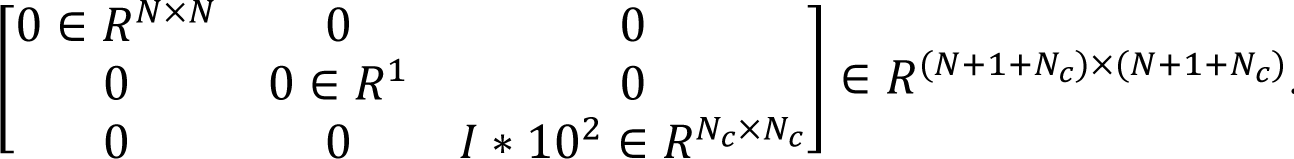. Thus, the final cursor state error is penalized, and the magnitude of the input to the neural population 𝑢_𝑡_ is penalized (with setting 𝑅 as non-zero). Because the magnitude of the input to neural activity is penalized, the controller sends the minimal input to the neural population to produce task behavior. We defined our cost function so that the cursor state during movement before the final cursor state is not penalized, and the neural state is never penalized.

The optimal control sequence (𝑢_𝑡_, 𝑡 = 0, 1, … 𝑇 − 1) is given by 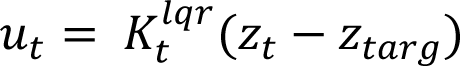 where feedback gain matrices 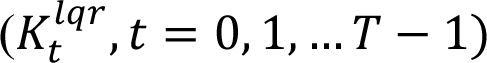 are computed iteratively solving the dynamic Ricatti equation backwards in time. We note that we computed the LQR solution for 𝑢_𝑡_ using the dynamics of state error *z_t_* − *z_targ_*, and that the dynamics of state error for non-zero target states are affine rather than strictly linear.

##### OFC for center-out task

Center-out task simulations were run with the initial cursor position in the center of the workspace at 𝑐_0_ = [0, 0, 0, 0, 1] and the target cursor state at [*target_x_,target_y_,vel_x_* = *vel_y_* = 0, 1]^*T*^. Targets were positioned 10cm away from the origin (same target arrangement as Monkey G). Target cursor velocity was set to zero to enforce that the cursor should stop at the desired target location.

Exact decoder parameters from Monkey G and linearized decoder parameters from Monkey J were used (𝐹, 𝐾) in simulations. The invariant neural dynamics model parameters (𝐴, 𝑏) were varied depending on the simulated experiment (see below). The horizon for each trial to hit its target state was set to be 𝑇 = 40 (corresponding to 4 seconds based on the BMI’s timebin of 100ms). Constraining each trial to be equal length facilitated comparison of performance across different simulation experiments. We verified that all of our simulated trials completed their tasks successfully.

##### OFC for obstacle-avoidance using a heuristic

Obstacle-avoidance task simulations were performed with the same initial and target cursor states as the center-out task, except that the cursor circumvented the obstacle to reach the target in both clockwise and counterclockwise movements. We used a heuristic strategy to direct cursor movements around the obstacle; we defined a waypoint as an intermediate state the cursor had to reach enroute to the final target. The heuristic solution performs optimal control from the start position to the waypoint, and then optimal control from the waypoint to the final target. Importantly, this solution minimizes the amount of input needed to accomplish these goals. We used a heuristic solution because the linear control problem of going from the initial cursor state to the final target cursor state with the constraint of avoiding an obstacle is not a convex optimization problem.

Concretely, for the first segment of the movement, a controller with a horizon T=20 directed the cursor to the waypoint, and then a controller with horizon T=20 directed the cursor from the waypoint to the final target (such that the trial length was matched to the center-out task simulation with T=40).

The waypoint was defined relative to the obstacle position as follows. First the vector between the center target and the obstacle position was determined (*v_obs,center_*). The *v_obs,center_* was then rotated either +90 degrees or -90 degrees corresponding to clockwise and counterclockwise movements. The waypoint position was a 6cm distance in the direction of the rotated vector, from the obstacle center. Finally, the desired velocity vector of the intermediate target was set to be in the direction of *v_obs,center_*, with a magnitude of 10 cm/s, so that the cursor would be moving in a direction consistent with reaching its final target in the second segment of the movement after the waypoint was reached.

To compute the input 𝑢_𝑡_ to execute these movements, we defined the state error at each time 𝑡 as *z_error_* = *Z_targ_* − *z_t_*, where *z_targ_* was the waypoint for the first half of the movement, and *z_targ_* was the final target for the second half of the movement. The linear quadratic regulator feedback gain 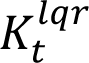 matrices were computed on the appropriate state error dynamics with the shortened horizon T=20.

##### “Full Dynamics Model” Simulation

Simulations of the “Full Dynamics Model” consisted of OFC with the invariant dynamics parameters (𝐴, 𝑏) that were fit on experimentally-recorded neural activity from each subject and session (see “Invariant dynamics models” below, under “Quantification and Statistical Analysis”). 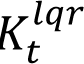 was computed using these experimentally-observed (𝐴, 𝑏) parameters. The initial state of neural activity (i.e. 𝑥_𝑡_ at t=0) was set to the fixed point of the dynamics.

##### “No Dynamics Model” Simulation

Simulations of the “No Dynamics Model” consisted of OFC with invariant dynamics parameter 𝐴 set to zero (𝐴 = 0). The experimentally-observed offset 𝑏 was still used from each subject and session. 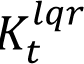 was computed using 𝐴 = 0 and the experimentally-observed 𝑏, and thus it was different than in the “Full Dynamics Model.” The initial state of neural activity (i.e. 𝑥_𝑡_ at t=0) was set to offset 𝑏, the fixed point of dynamics with 𝐴 = 0.

##### “Decoder-null Dynamics Model” Simulation

Simulations of the “Decoder-null Dynamics Model” consisted of OFC with the experimentally-observed invariant dynamics parameters (𝐴, 𝑏) that were restricted to the decoder-null space, i.e. each invariant dynamics model was fit only on the projection of neural activity into the decoder-null space (see “Invariant dynamics models” under “Quantification and Statistical Analysis”). 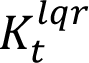 was computed using these experimentally-observed decoder-null (𝐴, 𝑏) parameters, and thus it was different than in the “Full Dynamics Model.” The initial state of neural activity (i.e. 𝑥_𝑡_ at t=0) was set to the fixed point of the decoder-null invariant dynamics.

The “Decoder-null Dynamics Model” was compared to its own “No Dynamics Model”, which consisted of OFC with 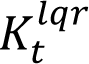 computed using 𝐴 = 0 and the experimentally-observed decoder-null offset 𝑏 for each subject and session, and thus it was different than in the previously defined models. The initial state of neural activity (i.e. 𝑥_𝑡_ at t=0) was set to the decoder-null offset 𝑏, the fixed point of dynamics with 𝐴 = 0.

##### Noise

In our OFC model, movement errors arise due to noise in the neural activity, and subsequent neural activity issues commands based on feedback to correct these errors. We used two considerations to choose the noise level for neural activity. First, we sought to add a level of neural noise that was comparable to the neural “signal” needed to perform control in the absence of noise. Second, we wanted to add the same level of noise to the dynamics model (either “Full Dynamics Model” or “Decoder-null Dynamics Model”) and the corresponding “No Dynamics Model,” in order to facilitate comparison.

Thus, we first simulated the “No Dynamics Model” without noise for a single trial for each of 24 conditions, and we calculated 𝑅, the average variance of a neuron across time and trials.

Then for our noisy simulations of the “No Dynamics Model” and the corresponding dynamics models, Gaussian noise with zero mean and fixed variance 𝑅 was added to each neuron at each timestep: 𝑥_𝑡_ = 𝐴𝑥_𝑡−1_ + 𝐵𝑢_𝑡−1_ + 𝜎_𝑡−1_, where 𝜎_𝑡_∼𝑁(0, 𝑅𝐼𝐼). Thus, the overall level of added noise (the sum of noise variance over neurons) matched the overall level of signal in the noiseless No Dynamics Model simulation (sum of activity variance over neurons).

We note that our main findings (Fig. 6CD, 6GH) held even with different noise levels.

#### QUANTIFICATION AND STATISTICAL ANALYSIS

##### Command discretization for analysis

We sought to analyze the occurrence of the same command across different movements. Commands on individual time points were analyzed as the same command if they fell within the same discretized bin of continuous-valued, two-dimensional command space. All commands from rewarded trials in a given experimental session (including both tasks) were aggregated and discretized into 32 bins. Individual commands were assigned to one of 8 angular bins (bin edges were 22.5, 67.5, 112.5, 157.5, 202.5, 247.5, 292.5, and 337.5 degrees) and one of four magnitude bins. Angular bins were selected such that the straight line from the center to each of the center-out targets bisected each of the angular bins as has been done in previous work^50^ (Fig. S1A). Magnitude bin edges were selected as the 23.75^th^, 47.5^th^, 71.25^th^, and 95^th^ percentile of the distribution of command magnitudes for that experimental session. Commands falling between the 95^th^ and 100^th^ percentile of magnitude were not analyzed to prevent very infrequent noisy observations from skewing the bin edges for command magnitude.

##### Conditions that used a command regularly

For each session, the number of times each of the 32 (discretized) commands was used in a given condition was tabulated. If the command was used >= 15 times for that condition within a given session pooling across trials, that condition was counted as using the command regularly and was used in all analyses involving (command, condition) tuples. Commands that were used < 15 times were not used in analysis involving (command, condition) tuples. We note that the main results of the study were not affected by this particular selection. Typically, an individual command is used regularly in 5-10 conditions (distribution shown in Fig. S1A).

#### Cursor and command trajectory visualization

##### Cursor position subtrajectories

To visualize the cursor position trajectories locally around the occurrence of a given command for each condition, we computed the average position “subtrajectory,” which we define as the average trajectory in a window locked to the occurrence of the given command. For each condition, cursor positions from successful trials were aggregated. Cursor position subtrajectories shown in Fig. 1F are from representative session 0 from Monkey G. A matrix of x-axis and y-axis position trajectories was formed by extracting a window of -500ms to 500ms (5 previous samples plus 5 proceeding samples) around each occurrence of the given command in a given condition (total of N_com-cond_ occurrences, yielding a 2 x 11 x N_com-cond_ matrix). Averaging over the N_com-cond_ observations yielded a condition-specific command-locked average position subtrajectory (size: 2 x 11) for each condition. If a command fell in the first 500ms or last 500ms of a trial, its occurrence was not included in the subtrajectory calculation. The position subtrajectories were translated such that the occurrence of the given command was set to (0, 0) in the 2D workspace (Fig. 1F *right*, Fig. S1C *middle*).

##### Command subtrajectories

To visualize trajectories of commands around the occurrence of a given command for each condition (Fig. 1G, *right*), we followed the same procedure as described above for cursor position subtrajectories to tabulate a 2 x 11 x N_com-cond_ matrix but with x-axis and y-axis commands instead of positions. We note that this matrix consisted of the continuous, two-dimensional velocity values of the commands. Averaging over the N_com-cond_ observations yielded the average condition-specific command subtrajectory (size: 2 x 11 array), as shown in Fig. 1F *left* for example conditions.

#### Matching the condition-pooled distribution

In many analyses, data (e.g. neural activity or a command-locked cursor trajectory) associated with a command and a specific condition is compared to data that pools across conditions for that same command (Figs. 3-5). The distribution of the precise continuous value of the command within the command’s bin may systematically differ between condition-specific and condition-pooled datasets, which we refer to as ‘within-command-bin differences.’ To ensure within-command-bin differences are not the source of significant differences between condition-specific and condition-pooled data associated with a command, we developed a procedure to subselect observations of condition-pooled commands so that the mean of the condition-pooled command distribution is matched to the mean of the condition-specific command distribution. This procedure ensures that any differences between the condition-specific quantity and condition-pooled quantity are not due to ‘within-command-bin differences’. This procedure is performed on all analyses comparing condition-specific data to a condition-pooled distribution of data. The matching procedure is as follows:

1. From the condition-specific distribution, compute the command mean 𝜇_*com-cond*_ (size: 2×1) and standard deviation 𝜇_*com-cond*_ (size: 2×1).
2. Compute the deviation of each continuous-valued command observation in the condition-pooled distribution from the condition-specific distribution.
  a. Use the condition-specific distribution’s parameters to z-score the condition-pooled distribution’s continuous-valued command observations by subtracting 𝜇_*com-cond*_ and dividing by 𝜇_*com-cond*_.
  b. Compute the deviation of condition-pooled observations from the condition-specific distribution as the L2-norm of the z-scored value
  c. For indices in the condition-pooled distribution that correspond to data in the condition-specific distribution, over-write the L2-norm of the z-scored values with zeros. This step prevents the condition-pooled distribution from dropping datapoints that are in the condition-specific data, thereby ensuring the condition-pooled distribution contains the condition-specific data.
3. Remove the 5% of condition-pooled observations with the largest deviations
4. Use a Student’s t-test to assess if the remaining observations in the condition-pooled distribution are significantly different than the condition-specific distribution for the first and second dimension of the command (two p-values)
5. If both p-values are > 0.05, then the procedure is complete and the remaining observations in the condition-pooled distribution are considered the “command-matched condition-pooled distribution” for a specific command and condition.
6. If either or both p-values are < 0.05, return to step 3 and repeat.

If the condition-pooled distribution cannot be matched to the condition-specific distribution such that the size of the condition-pooled distribution is larger than the condition-specific distribution, the particular (command, condition) will not be included in the analysis.

#### Comparing command subtrajectories

To assess whether a command is used within significantly different command subtrajectories in different conditions (Fig. S1 DE), the following analysis is performed for conditions that have sufficient occurrences of the command (>=15):

1. The condition-specific average command subtrajectory is computed by averaging over N_com-cond_ single-trial command subtrajectories for the condition, as defined above in “*Visualization of command subtrajectories*”.
2. The condition-pooled average command subtrajectory is computed: all the single-trial command subtrajectories (N_com_) are pooled across trials from all conditions that use the given command regularly (command occurs >= 15 times in a session) to create a condition-pooled distribution of single-trial command subtrajectories (a 2 x 11 x N_com_ matrix), which is then averaged to yield the condition-pooled average command subtrajectory (a 2 x 11 matrix).
3. In order to test whether condition-specific average command subtrajectories were significantly different from the condition-pooled average command subtrajectory, a distribution of subtrajectories was created by subsampling the condition-pooled distribution to assess expected variation in subtrajectories due to limited data. Specifically, N_com-cond_ single-trial command subtrajectories were sampled from a condition-pooled distribution of command subtrajectories that was command-matched to the specific condition (see above, “Matching the condition-pooled distribution”). These N_com-cond_ samples were then averaged to create a single subtrajectory, representing a plausible condition-specific average subtrajectory under the view that the condition-specific subtrajectories are just subsamples of the condition-pooled subtrajectories. This procedure was repeated 1000 times and used to construct a bootstrapped distribution of 1000 command subtrajectories.
4. This distribution was then used to test whether condition-specific subtrajectories deviated from the condition-pooled subtrajectory more than would be expected by subsampling and averaging the condition-pooled subtrajectory distribution. Specifically, the true condition-specific command subtrajectory distance from the condition-pooled command subtrajectory was computed (L2-norm between condition-specific 2×11 subtrajectory and condition-pooled 2×11 subtrajectory) and compared to the bootstrapped distribution of distances: (L2-norm between each of the 1000 subsampled averaged 2×11 command subtrajectories and the condition-pooled 2×11 command subtrajectory). A p-value for each condition-specific command subtrajectory distance was then derived.

The same analysis is also performed using only the next command following a given command (Fig. S1 E).

#### Behavior-preserving shuffle of activity

We shuffled neural activity in a manner that preserved behavior as a control for comparison against the hypothesis that neural activity follows invariant dynamics beyond the structure of behavior. Shuffled datasets preserved the timeseries of discretized commands but shuffled the neural activity that issues these commands. In order to create a shuffle for each animal on each session, all timebins from all trials from all conditions were collated. The continuous-valued command at each timebin was labeled with its discretized command bin. For each of the 32 discretized command bins, all timebins corresponding to a particular discretized command bin were identified. The neural activity in these identified timebins was then randomly permuted. A complete shuffled dataset was constructed by performing this random permutation for all discretized command bins. This full procedure was repeated 1000 times to yield 1000 shuffled datasets.

#### Analysis of activity issuing a given command

##### Condition-specific neural activity distances

For each session, (command, condition) tuples with >= 15 observations were analyzed. For each of these (command, condition) tuples, we analyzed the distance between condition-specific average activity and condition-pooled average activity, both for individual neurons and for the population’s activity vector (Fig. 3B-E). Analysis of individual neurons for a given (command, condition) tuple, given 𝑁 neurons:

1. Compute the condition-specific average neural activity (𝜇_*com-cond*_ ∈ 𝑅^𝑁^) as the average neural activity over all observations of the command in the condition.
2. Compute the condition-pooled average activity (𝜇_*com-cond*_ ∈ 𝑅^𝑁^) as the average neural activity over observations of the command pooling across conditions. The command-matching procedure is used to form the condition-pooled dataset to account for within-command-bin differences (see “Matching the condition-pooled distribution” above).
3. Compute the absolute value of the difference between the condition-specific and condition-pooled averages: 𝑑𝜇_*com*−*cond*_ = 𝑅𝑏𝑠(𝜇_*com*−*cond*_-𝜇_*com−pool*_) ∈ 𝑅^𝑁^.
4. Repeat steps 1-3 for each shuffled dataset *i*, yielding 𝑑𝜇_𝑒ℎ𝑢−𝑖−*com*−*cond*_ for 𝑑 = 1: 1000.
5. For each neuron *j,* compare 𝑑𝜇_*com*−*cond*_(𝑗) to the distribution of 𝑑𝜇_𝑒ℎ𝑢−𝑖−*com*−*cond*_(𝑗) for i = 1:1000. Distances greater than the 95^th^ percentile of the shuffled distribution are deemed to have significantly different neuron *j* activity for a command-condition. Analysis of population activity for a given (command, condition) tuple:

To compute population distances, one extra step was performed. We sought to ensure that the distances we calculated were not trivially due to “within-bin differences” between the condition-specific and condition-pooled distributions. The first step to ensure this was described above in “Matching the condition-pooled distribution”. The second step was to only compute distances in the dimensions of neural activity that are null to the decoder and do not affect the composition of the command. Thus, any subtle remaining differences in the distribution of commands would not influence population distances.

To compute distances in the dimensions of neural activity null to the decoder, we computed an orthonormal basis of the null space of decoder matrix 𝐾 ∈ 𝑅^2𝑥𝑁^ using scipy.linalg.null_space, yielding 𝑉_𝑥𝑢𝑥_ ∈ 𝑅^𝑁𝑥𝑁−2^. The columns of 𝑉 correspond to basis vectors spanning the 𝑁 − 2 dimensional null space. Using 𝑉_𝑥𝑢𝑥_ we computed: 𝜇_*com*−*cond*−𝑥𝑢𝑥_ = 𝑉_𝑥𝑢𝑥_′ ∗ 𝜇_*com*−*cond*_ and 𝜇_*com*−𝑝𝑜𝑜𝑥−𝑥𝑢𝑥_ = 𝑉_𝑥𝑢𝑥_′ ∗ 𝜇_*com*−*pool*_. We then calculated the population distance metric (L2-norm), normalized by the square-root of the number of neurons: 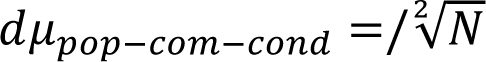, 𝑑𝜇_𝑝𝑜𝑝−*com*−*cond*_ ∈ 𝑅^1^. In step 5, the single value 𝑑𝜇_𝑝𝑜𝑝−*com*−*cond*_ is compared to the distribution of 𝑑𝜇_𝑒ℎ𝑢−𝑖−𝑝𝑜𝑝−*com*−*cond*_for i = 1:1000 to derive a p-value for each (command, condition) tuple. The fraction of (command, condition) tuples with population activity distances greater than the 95^th^ percentile of the shuffle data (i.e. significant) is reported in Fig. 3E.

For visualization of distances relative to the shuffle distribution (Fig. 3B-D), we divided the observed population distance for each (command, condition) tuple by the mean of the corresponding shuffle distribution. With this normalization, we can visualize the spread of the shuffle distribution (Fig. 3B, *right*) and we can interpret a normalized distance of 1 as the expected distance according to the shuffle distribution.

##### Activity distances pooling over conditions

To test whether condition-specific neural activity significantly deviated from condition-pooled neural activity for a given command (Fig. 3E, *middle*), we aggregated the distance between condition-specific and condition-pooled average activity over all 𝑁𝑐𝑝𝑅𝑑 conditions in which the command was used (>= 15 occurrences of the command in a condition) . An aggregate command distance is computed: 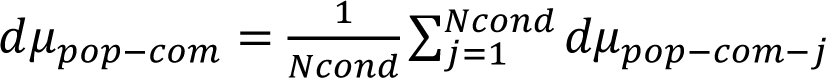, and an aggregate shuffle distribution is computed: 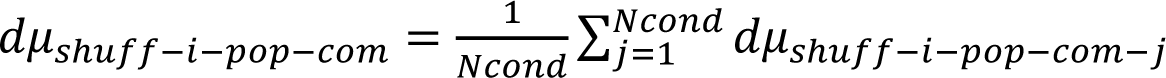. Then *d*_βpop-com_ is compared to the distribution of 𝑑𝜇_𝑒ℎ𝑢−𝑖−𝑝𝑜𝑝−*com*_ for 𝑑 = 1: 1000 to derive a p-value for each command. The fraction of commands with significant population activity distances is reported in Fig. 3E, *middle*.

##### Single neuron distances

To test whether an individual neuron’s condition-specific activity deviated from condition-pooled activity (Fig. 3E *right*), we aggregated the distances between condition-specific and condition-pooled average activity over the 𝐶 (command, condition) tuples with at least 15 observations. The aggregated distance for neuron 𝑅 was computed: 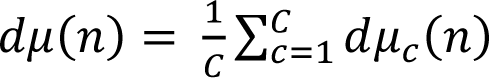 where 𝑑𝜇_𝑐_(𝑅) is the condition-specific absolute difference for the 𝑅th neuron and 𝑐th (command, condition) tuple. Then 𝑑𝜇(𝑅) was compared to the distribution of the aggregated shuffle: 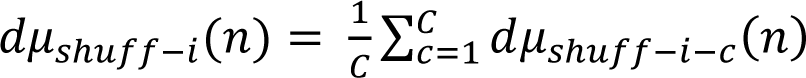 to derive a p-value for each neuron. The fraction of neurons with significant activity distances (p-value<0.05) is reported in Fig. 3E *right*.

##### Neural activity distances summary

Single neuron activity distances reported in Fig. S2B (*left)* are for all (command, condition, neuron) tuples that had at least 15 observations. We report distances as a z-score of shuffle distribution: 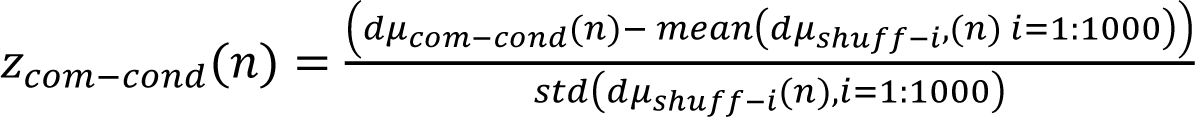.

Single neuron activity distances reported in (Fig. S2B *center, right*) are for (command, condition, neuron) tuples that significantly deviated from shuffle. We report raw distances in neuron activity as 𝑑𝜇 *com*−*cond* (𝑅) (Fig. S2B, *center*), and fraction distances as 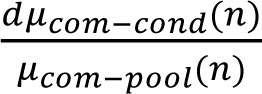 (Fig. S2B, *right*).

Population activity distances reported in Fig. 3BCD and Fig. S2C *left* are for all (command, condition) tuples. We report distances in population activity as a fraction of shuffle mean: 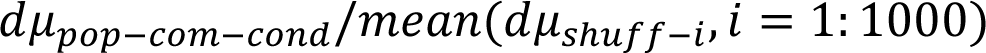 (Fig. 3BCD), and as a z-score of shuffle distribution: 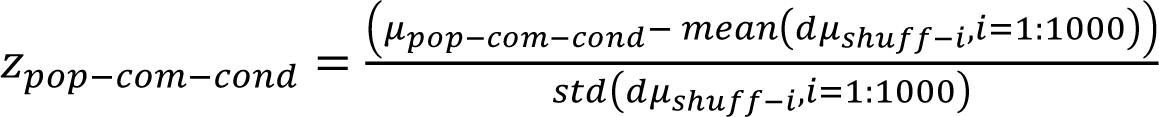 (Fig. S2C *left*).

Population activity distances reported in Fig. S2C (*center, right)* are for (command, condition) tuples that significantly deviated from shuffle. We report distances in population activity as a fraction of shuffle mean 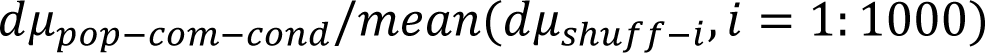 (Fig. S2C, *center*) and fraction of condition-pooled activity as 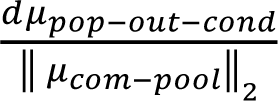 (Fig. S2C, *right*).

#### Invariant dynamics models

In order to test whether invariant dynamics predicts the different neural activity patterns issuing the same command for different conditions, a linear model was fit for each experimental session on training data of neural activity from all conditions and assessed on held-out test data. Neural activity at time *t,* 𝑥_𝑡_, was modeled as a linear function of 𝑥_𝑡−1_:

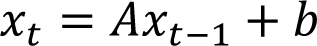

Here 𝐴 ∈ 𝑅^𝑁𝑥𝑁^ modeled invariant dynamics and 𝑏 ∈ 𝑅^𝑁^ was an offset vector that allowed the model to identify non-zero fixed points of neural dynamics. Ridge regression was used to estimate the 𝐴 and 𝑏 parameters. Prior to any training or testing, data was collated such that all neural activity in bins from t=2:Ttrl in all rewarded trials were paired with neural activity from t=1:(Ttrl-1), where Ttrl is the number of time samples in a trial.

##### Estimation of Ridge Parameter

For each experimental session, data collated from all conditions was randomly split into 5 sections, and a Ridge model (sklearn.linear_model.Ridge) with a ridge parameter varying from 2.5×10^-5^ to 10^6^ was trained using 4 of the 5 sections and tested on the remaining test section. Test sections were rotated, yielding five estimates of the coefficient of determination (R^2^) for each ridge parameter. The ridge parameter yielding the highest cross-validated mean R^2^ was selected for each experimental session. Ridge regression was used primarily due to a subset of sessions with a very high number of units (148 and 151 units), thus a high number of parameters needed to be estimated for the 𝐴 matrix. Without regularization, these parameters tended to extreme values, and the model generalized poorly.

##### Invariant dynamics model: fitting and testing

Once a ridge parameter for a given experimental session was identified, 𝐴, 𝑏 were again trained using 4/5 of the data. The remaining test data was predicted using the fit 𝐴, 𝑏. This procedure was repeated, rotating the training and testing data such that after five iterations, all data points in the experimental session had been in the test data section for one iteration of model-fitting. The predictions made on the held-out test data were collated together into a full dataset. Predictions were then analyzed in subsequent analyses.

##### Generalization of invariant dynamics

We assessed how well invariant dynamics generalized when certain categories of neural activity were not included in the training data. Invariant dynamics models were estimated after excluding neural activity in the following categories (Fig. 4C, Fig. S4, Fig 5CD):

1. Left-out Command: For each command (total of 32 command bins), training data sets were constructed leaving out neural activity that issued the command (Fig. 4C, Fig. S4, Fig. 5CE).
2. Left-out Condition: For each condition (consisting of target, task, and clockwise or counterclockwise movement for obstacle avoidance), training data sets were constructed leaving out neural activity for the given condition (Fig. 4C, Fig. S4, Fig. 5CE).
3. Left-out Command Angle: For each command angular bin (total of 8 angular bins), training data sets were constructed leaving out neural activity that issued commands in the given angular bin. This corresponds to leaving out neural activity for the 4 command bins that have the given angular bin but different magnitude bins (Fig. S4B, middle).
4. Left-out Command Magnitude: For each command magnitude bin (total of 4 magnitude bins), training data sets were constructed leaving out neural activity that issued commands of the given command magnitude. This corresponds to leaving out neural activity for the 8 command bins that have the given magnitude bin but different angle bins (Fig. S4B, right).
5. Left-out Classes of Conditions (Fig. S4G):
  a. vertical condition class consisting of conditions with targets located at 90 and 270 degrees for both tasks,
  b. horizontal condition class consisting of conditions with targets located at 0 and 180 degrees for both tasks,
  c. diagonal 1 condition class consisting of conditions with targets located at 45 and 215 degrees for both tasks, and
  d. diagonal 2 condition class consisting of conditions with targets located at 135 and 315 degrees for both tasks.

For each of the listed categories above, many dynamics models were computed – each one corresponding to the exclusion of one element of the category (i.e. one model per: command left-out, condition left-out, command angle left-out, command magnitude left-out, and class of conditions left-out). Each of the trained models was then used to predict the left-out data. Predictions were aggregated across all dynamics models resulting in a full dataset of predictions. The coefficient of determination (R^2^) of this predicted dataset reflected how well dynamics models could generalize to types of neural activity that were not observed during training. We note that Monkey J did not perform all conditions in the “diagonal 2” class, and so was not used in the analysis predicting excluded “diagonal 2” conditions.

##### Decoder-null dynamics model

As an additional comparison, we modeled invariant dynamics that lie only within the decoder-null space (the neural activity subspace that was orthogonal to the decoder such that variation of neural activity in this space has no effect on the decoder’s output, i.e. commands for movement).

Our approach was to project spiking activity into the decoder null space, and then fit invariant dynamics on the projected, decoder-null spiking activity. We first computed an orthonormal basis of the null space of decoder matrix 𝐾 ∈ 𝑅^2𝑥𝑁^ using scipy.linalg.null_space, yielding 𝑉_𝑥𝑢𝑥_ ∈ 𝑅^𝑁𝑥𝑁−2^. The columns of 𝑉 correspond to basis vectors spanning the 𝑁 − 2 dimensional null space. We then computed the projection matrix *P_null_* ϵ *R^NxN^* where *P_null_* = *V_null_V_null_^T^*. Spiking activity was then projected into the null space *x_t_^null^*, where *x_t_^null^* ϵ *R^Nx1^*.

Following the above procedure (see “*Estimation of Ridge Parameter”)*, a ridge regression parameter was selected using projected data *x_t_^null^*. Decoder-null dynamics model parameters *Anull, bnull* were then fit on 4/5 of the dataset and then tested on the remaining 1/5 of the *x_t_^null^* dataset. As before, the training/testing procedure was repeated 5 times such that all data points fell into the test dataset once. Predictions of test data from all five repetitions were collated into one full dataset of predictions. We note that the average of the decoder-space activity across the entire session 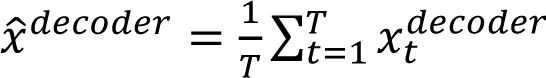, where 𝑇 is the number of bins in an entire session, was added to all predictions of decoder-null dynamics 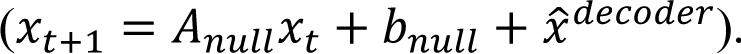.

##### Shuffle dynamics model

The invariant dynamics model was compared to a shuffle dynamics model fit on shuffled data (see “<u>Behavior-preserving shuffle of activity”</u> above). Following the above procedure (see “*Estimation of Ridge Parameter”)*, a ridge parameter was selected using shuffled data. Shuffle dynamics model parameters *Ashuffle, bshuffle* were then fit on 4/5 of the dataset using shuffled data and then tested on the remaining 1/5 of the dataset using original, unshuffled data.

#### Invariant dynamics model characterization

##### Dimensionality and eigenvalues

Once the linear invariant dynamics model’s parameters *A, b* were estimated, *A* was analyzed to assess which modes of dynamics^16^ were present (Fig. S3). The eigenvalues of *A* were computed. From each eigenvalue, an oscillation frequency and time decay value were computed using the following equations:

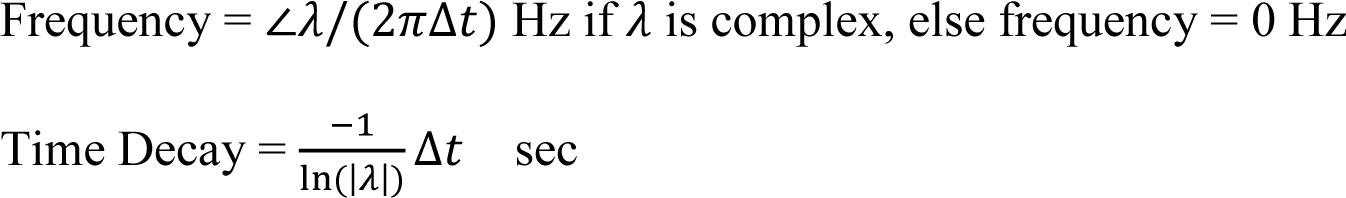

Modes of dynamics contributing substantially to predicting future neural variance will have time decays greater than the BMI decoder’s binsize (here, 100ms). 2-4 such dimensions of dynamics were found across sessions and subjects (Fig. S3).

#### Invariant dynamics model predictions

##### Predicting next neural activity: 𝑥_𝑡+1_| 𝑥_𝑡_, 𝐴, 𝑏

In Fig. 5C, we predict next activity 𝑥_𝑡+1_ based on current activity 𝑥_𝑡_ by taking the expected value according to our model: 𝐸(𝑥_𝑡+1_|𝑥_𝑡_, 𝐴, 𝑏) = 𝐴𝑥_𝑡_ + 𝑏.

In Fig. 5D, we evaluated this prediction for individual dimensions of neural activity. We projected the prediction of 𝑥_𝑡+1_ onto each eigenvector of the dynamics model 𝐴 matrix and evaluated how well that dimension was predicted (via coefficient of determination).

In Fig. S3E, G, we evaluated this prediction across time from the start of trial. The magnitude (i.e. L2 norm) of the model residual ‖𝑥_𝑡+1_ − 𝐴𝑥_𝑡_ + 𝑏‖_2_ (Fig. S3E) and the coefficient of determination (R^2^) (Fig. S3G) are plotted for each time point from trial start, evaluated on held-out test data pooling across trials.

##### Predicting next command: command_𝑡+1_| 𝑥_𝑡_, 𝐴, 𝑏, 𝐾

In Fig. 5E-H, we predict the next command command_𝑡+1_ based on current neural activity 𝑥_𝑡_ by taking its expected value according to our model: 𝐸(command_𝑡+1_ | 𝑥_𝑡_, 𝐴, 𝑏, 𝐾) = 𝐾(𝐴𝑥_𝑡_ + 𝑏), where the decoder matrix *K* maps between neural activity and the command. This amounts to first predicting next activity based on current activity as above 𝐸(𝑥_𝑡+1_|𝑥_𝑡_, 𝐴, 𝑏) = 𝐴𝑥_𝑡_+ 𝑏 and then applying decoder *K*.

##### Predicting activity issuing a given command

In Fig. 4C-G, we predict current activity 𝑥_𝑡_ not only with knowledge of previous activity 𝑥_𝑡−1_, but also with knowledge of the current command command_𝑡_ (𝑥_𝑡_| 𝑥_𝑡−1_, 𝐴, 𝑏, 𝐾, command_𝑡_). We modeled 𝑥_𝑡_ and 𝑥_𝑡−1_ as jointly Gaussian with our dynamics model, and command_𝑡_ is jointly Gaussian with them since command_𝑡_ = 𝐾𝑥_𝑡_. We modify our prediction of 𝑥_𝑡_ based on knowledge of command_𝑡_: 𝐸(𝑥_𝑡_|𝑥_𝑡−1_, 𝐴, 𝑏, 𝐾, command_𝑡_). Explicitly we conditioned on command_𝑡_, thereby ensuring that 𝐾 ∗ 𝐸(𝑥_𝑡_|𝑥_𝑡−1_, 𝐴, 𝑏, 𝐾, command_𝑡_) = command_𝑡_. To do this we wrote the joint distribution of 𝑥_𝑡_ and command_𝑡_:

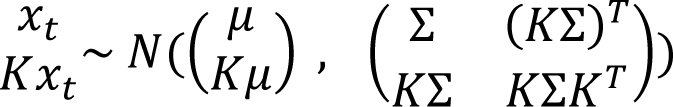

where 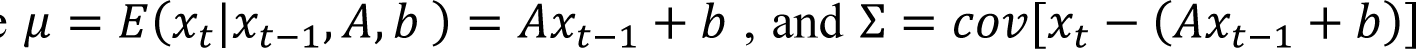 is the covariance of the noise in the dynamics model. Then, the multivariate Gaussian conditional distribution provides the solution to conditioning on command_𝑡_:

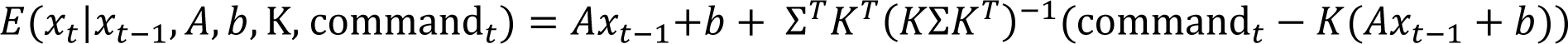

This prediction constrains the prediction of 𝑥_𝑡_ to produce the given command command_𝑡_.

For these predictions, Σ is estimated following dynamics model fitting and set to the empirical error covariance between estimates of 𝐸(𝑥_𝑡_) = 𝐴𝑥_𝑡−1_ + 𝑏 and true 𝑥_𝑡_ in the training data.

##### Predicting current activity only with command

In Fig. 4C-E, as a comparison to the dynamics prediction (𝑥_𝑡_| 𝑥_𝑡−1_, 𝐴, 𝑏, 𝐾, command_𝑡_), we predict 𝑥_𝑡_ as its expected value (𝑥_𝑡_| 𝐾, command_𝑡_) based only on the command command_𝑡_ = 𝐾𝑥_𝑡_ it issues and the decoder matrix 𝐾. The same approach was used as above, except with empirical estimates of 𝜇, Σ corresponding to the mean and covariance of the neural data instead of using the neural dynamics model and 𝑥_𝑡−1_ to compute 𝜇, Σ.

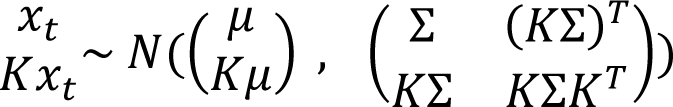

This formulation makes the prediction:

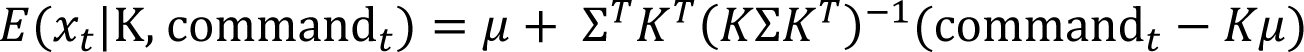

##### Comparing invariant dynamics to shuffle

For the above predictions, we evaluated if invariant dynamics models were more accurate than shuffle dynamics. A distribution of shuffle dynamics R^2^ values (coefficient of determination) was generated by computing one R^2^ value per shuffled dataset (see “<u>Behavior-preserving shuffle of activity”</u> above), where 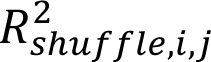 corresponds to the 𝑅^2^ for shuffle dataset 𝑑 on session 𝑗. For each session 𝑗, each invariant dynamics model was considered significant if its R^2^ was greater than 95% of shuffle R^2^ values. To aggregate over 𝑆 sessions, the R^2^ values for all 𝑆 sessions were averaged yielding one 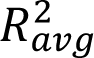 value. This averaged value was compared to a distribution of averaged shuffle R^2^ values. Specifically, for each shuffle 𝑑 (i=1:1000 shuffled dataset) an averaged R^2^ value was computed across all 𝑆 sessions: 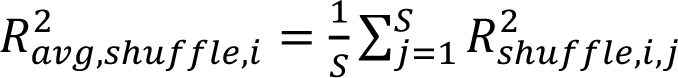, yielding a distribution of averaged shuffle R^2^ values.

##### Predicting condition-specific activity

The invariant dynamics model was used to predict the condition-specific average activity for a given command (𝜇_*com*−*cond*_, i.e. the average neural activity over all observations of the command in the condition, see “<u>Analysis of activity issuing a given command</u>” above) (Fig. 4D-G). The invariant dynamics model prediction 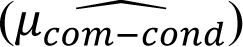 was computed as 𝐸(𝑥_𝑡_|𝑥_𝑡−1_, 𝐴, 𝑏, 𝐾, command_𝑡_) (see “*Predicting activity issuing a given command*” above) averaged over all observations of neural activity for the given command and condition.

To test if the invariant dynamics prediction was significantly more accurate than the shuffle dynamics model (i.e. the dynamics model fit on shuffled data, see “Shuffle dynamics model” above) prediction, we computed the error as the distance between true (𝜇_*com*−*cond*_) and predicted 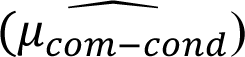 condition-specific average activity (single neuron error and population distance). Note that population distances for true and predicted activity were taken only in the dimensions null to the decoder (see “*Condition-specific neural activity deviation”*). The invariant dynamics model was deemed significantly more accurate than shuffle dynamics if the error was less than the 5^th^ percentile of the distribution of the errors from shuffle dynamics models. We reported the fraction of (command, condition) tuples that were individually significant relative to shuffle (Fig. 4G, left). We determined whether commands were individually significant relative to shuffle by analyzing the average population activity error across conditions (Fig 4G, middle). We determined whether neurons were individually significant relative to shuffle by analyzing the average single-neuron error over (command, condition) tuples (Fig 4G, right).

##### Predicting condition-specific component

The component of neural activity for a given command that was specific to a condition was calculated as 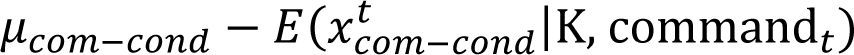, where 𝜇_*com*−*cond*_ is neural activity averaged over observations for the given command and condition, and 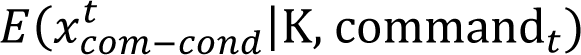 is the prediction of neural activity only given the command it issued, averaged over observations for the (command, condition) tuple (see “*Predicting current activity only with command”* above). Thus, 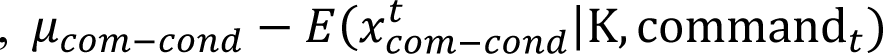 estimates the portion of neural activity that cannot be explained by just knowing the command issued.

We analyzed how well this condition-specific component could be predicted with invariant dynamics as: 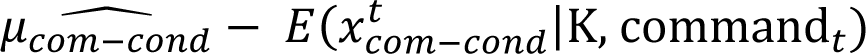 (see “*Predicting condition-specific activity*” above for calculation of 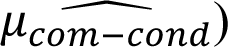. The variance of 𝜇_*com*−*cond*_ − 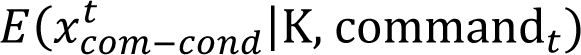 explained by 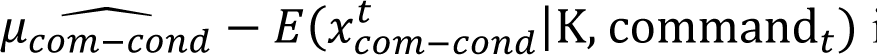 is reported *com*−*cond* in Fig. 4F.

##### Predicting condition-specific next command

For each (command, condition) tuple, the average “next command” command_*com*−*cond*_was calculated. For every observation of the given command in the given condition, we took the command at the time step immediately following the given command and averaged over observations. We then analyzed how well invariant dynamics predicted this average “next command” 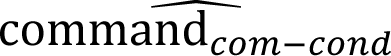, calculated as 𝐸(command_𝑡+1_ | 𝑥_𝑡_, 𝐴, 𝑏, 𝐾) averaged over all observations of neural activity 𝑥_𝑡_ for the given command and condition. The L2-norm of the difference 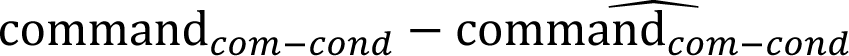 was computed and compared to the errors obtained from the shuffled-dynamics predictions. For each (command, condition) tuple, the dynamics-predicted “next command” was deemed significantly more accurate than shuffle dynamics if the error was less than the 5^th^ percentile of the distribution of the errors of the shuffled-dynamics predictions (Fig. 5F, *left*). Commands were determined to be individually significant if the error averaged over conditions was significantly less than the shuffled-dynamics error averaged over conditions (Fig. 5F, *right*).

##### Analysis of predicted command angle

We sought to further analyze whether invariant dynamics predicted the transition from a given command to different “next commands” in different movements. Thus, we calculated two additional metrics on the direction of the predicted “next command”, i.e. the angle of the predicted “next command” 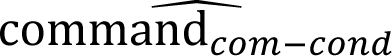 with respect to the condition-pooled “next command” command_*com*−*pool*_ (the average “next command” following a given command when pooling over conditions).

First, we predicted whether a condition’s “next command” would rotate clockwise or counterclockwise relative to the condition-pooled “next command.” Specifically, we calculated whether the sign of the cross-product between 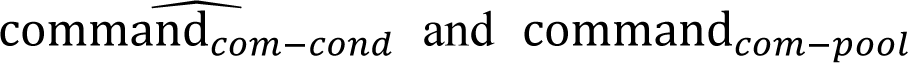 matched the sign of the cross-product between command_*com*−*cond*_ and command_*com*−*pool*_. The fraction of (command, conditions) that were correctly predicted (clockwise vs counterclockwise) was compared to the fraction of (command, condition) tuples correctly predicted in the shuffle distribution (Fig. 5H, *left*).

Second, we calculated the absolute error of the angle between the predicted “next command” and the condition-pooled “next command” for each (command, condition) tuple:

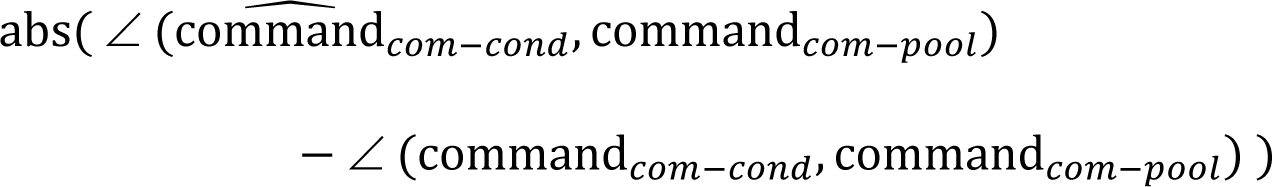

Explicitly, for each (command, condition) tuple, we calculated the absolute difference between two angles: 1) the angle between the predicted “next command” and the condition-pooled “next command” and 2) the angle between the true “next command” and the condition-pooled “next command”. These errors were then compared to the shuffle distribution (Fig. 5H, *right*).

#### Estimation of behavior-encoding models

To compare invariant dynamics models to models in which neural activity encodes behavioral variables in addition to the command, we fit a series of behavior-encoding models (Fig. S5). Regressors included cursor state (position, velocity), target position (x,y postion in cursor workspace), and a categorical variable encoding target number (0-7) and task (“center-out”, “clockwise obstacle-avoidance”, or “counter-clockwise obstacle-avoidance”).

Models were fit using Ridge regression following the same procedure described above (see “*Estimation of Ridge Parameter*”) was followed with one additional step: prior to estimating the ridge parameter or fitting the regression, variables were z-scored. Without z-scoring, ridge regression may favor giving explanatory power to the variables with larger variances, since they would require smaller weights which ridge regression prefers. Then, as above, models were fit using 4/5 of the data and then used to predict the held-out 1/5 of data. After 5 rotations of training and testing data, a full predicted dataset was collated.

We then tested whether invariant neural dynamics improved the prediction of neural activity beyond behavior-encoding. The coefficient of determination (R^2^) of the model containing all regressors except previous neural activity was compared to the R^2^ of the model containing all regressors plus previous neural activity (Fig. S5B) using a paired Student’s t-test where session was paired. One test was done for each monkey.

#### Analysis between pairs of conditions

We sought to assess whether the invariant dynamics model predicted the relationship between pairs of conditions for neural activity and behavior (Fig. S6).

##### Average neural activity for a given command

The invariant dynamics model was used to predict the distance between average neural activity patterns for the same command across pairs of conditions. Concretely, the predicted distance was simply the distance between the predicted neural activity pattern for condition 1 and for condition 2. The correlation between the true distance and the predicted distance was reported for individual neurons (Fig. S6AC) and population activity (Fig. S6BD). The Wald test (implemented in scipy.stats.linregress) was used to assess the significance of the correlations on single sessions. To assess significance pooled over sessions, data points (true distances vs. dynamics model predicted distances) were aggregated across sessions and assessed for significance.

##### Average next command

The invariant dynamics model was used to predict the distance between “next commands” for the same given command across pairs of conditions. Concretely, the predicted distance was simply the distance between the predicted “next command” for condition 1 and for condition 2. The correlation between the true distance and the predicted distance was reported (Fig. S6JK). As above, the Wald test was used to assess significance of correlations on single sessions and over pooled sessions.

##### Correlating neural distance with behavior

We asked whether neural activity for a given command was more similar across conditions with more similar command subtrajectories (see “*Command subtrajectories”*) (Fig. S6E), and whether invariant dynamics predict this. Specifically, we analyzed whether the distance between average neural activity across two conditions for a given command correlated to the distance between command subtrajectories for the same two conditions (Fig. S6, F *top,* GH *left*). Further, we analyzed whether invariant dynamics predicted this correlation (Fig. S6, F *bottom,* GH *right*). For every command (that was used in more than five conditions) and pair of conditions that used the command (>=15 observations in each condition in the pair), 1) the distances between condition-specific average activity were computed and 2) distances between command subtrajectories were computed. The neural activity distances were correlated with the command subtrajectory distances (Fig. S6, F *top,* GH *left*) . To assess whether invariant dynamics made predictions that maintained this structure, we performed that same analysis with distances between dynamics-predicted condition-specific average activity across pairs of conditions (Fig. S6, F *bottom,* GH *right*).

We assessed the significance of the relationship using a linear mixed effects (LME) model (statsmodels.formula.api.mixedlm). The LME modeled command as a random effect because the exact parameters of the increasing linear relationship between command subtrajectories and population activity may vary depending on command. Individual sessions were assessed for significance. To assess significance across sessions, data points were aggregated over sessions, and the LME model used command and session ID as random effects.

#### Analysis of Optimal Feedback Control Models

##### Input magnitude

For each simulated trial, we computed the magnitude of input to the neural population as the L2 norm of the input matrix 𝑢_𝑡_ ∈ 𝑅^𝑁×𝑇^ (where 𝑁 is the number of neurons and 𝑇 = 40 was the horizon and thus movement length). For each of the 24 conditions, we calculated the average input magnitude over the 20 trials. We compared the magnitude of input used by the Invariant Dynamics Model and the No Dynamics Model, where the Invariant Dynamics Model was either the Full Dynamics Model (Fig. 6C) or the Decoder-Null Dynamics Model (Fig. 6D). We analyzed each individual session with a paired Wilcoxon signed-rank test, where each pair within a session consisted of one condition (24 conditions total). We aggregated across sessions for each subject using a linear mixed effect (LME) model between input magnitude and model category (Invariant Dynamics Model or No Dynamics Model), with session modeled as a random effect.

##### Simulated activity issuing a given command

In the OFC simulations, we sought to verify if different neural activity patterns were used to issue the same command across different conditions, applying analyses that we used on experimental neural data to the OFC simulations. As above, we defined discretized command bins (see <u>“Command discretization for analysis”</u>) and calculated the average neural activity for each (command, condition) tuple. For (command, condition) tuples with >=15 observations (example shown in Fig. 6E), we computed the distance between condition-specific average activity and condition-pooled average activity by subtracting the activity, projecting into the decoder-null space, taking the L2 norm, and normalizing by the square root of the number of neurons, as in the experimental data analysis (see “<u>Analysis of activity issuing a given command</u>”).

We analyzed the distance between condition-specific average activity and condition-pooled average activity for a given command, comparing each model to its own shuffle distribution (see “<u>Behavior-preserving shuffle of activity</u>”) (Fig. 6GH). Concretely, for each simulated session, we calculated the mean of the shuffle distribution of distances for each (command, condition) tuple and compared these shuffle means (one per (command, condition) tuple) to the observed distances from the simulations. We analyzed individual sessions with a Mann-Whitney U test. We aggregated across sessions for each subject with a LME model between activity distance and data source (OFC Simulation vs shuffle), with session modeled as a random effect. For visualization of distances relative to the shuffle distribution (Fig. 6F-H), we divided the observed distance for each (command, condition) tuple by the mean of the corresponding shuffle distribution (same as in Fig. 3B-D).

#### Statistics Summary

In many analyses, we assessed whether a quantity calculated for a specific condition was significantly larger than expected from the distribution of the quantity due to subsampling the condition-pooled distribution. A p-value was computed by comparing the condition-specific quantity to the distribution of the quantity computed from subsampling the condition-pooled distribution. The “behavior-preserving shuffle of activity” and “<u>matching the condition-pooled distribution</u>” (see above) were used to construct the condition-pooled distribution.

The following is a summary of these analyses:

- Fig. S1D, Quantity: distance between condition-specific average command subtrajectory and condition-pooled average command subtrajectory, P-value: computed using behavior-preserving shuffle.
- Fig. S1E, Quantity: distance between condition-specific average next command and the condition-pooled average next command, P-value: computed using behavior-preserving shuffle.
- Fig. 3B *left*, 3E *right*: Quantity: for a given command, distance between condition-specific average activity for a neuron and condition-pooled average activity for a neuron, P-value: behavior-preserving shuffle.
- Fig. 3B *right*, 3D, 3E *left, middle*: Quantity: for a given command, distance between condition-specific average population activity and condition-pooled average population activity, P-value: behavior-preserving shuffle.
- Fig. 4G *right:* Quantity: for a given command, error between the invariant dynamics’ prediction of condition-specific average activity for a neuron and the true condition-specific average activity for the neuron. P-value: distribution of prediction errors from shuffle dynamics (models fit on behavior-preserving shuffle and that made predictions using unshuffled data).
- Fig. 4G *left, middle:* Quantity: for a given command, error between the invariant dynamics’ prediction of condition-specific average population activity and the true condition-specific average population activity. P-value: distribution of prediction errors from shuffle dynamics (models fit on behavior-preserving shuffle and that made predictions using unshuffled data).
- Fig. 5F: Quantity: for a given command, error between the invariant dynamics’ prediction of condition-specific average next command and true condition-specific average next command. P-value: distribution of prediction errors from shuffle dynamics (models fit on behavior-preserving shuffle and that made predictions using unshuffled data).

In the above analyses, we also assessed the fraction of condition-specific quantities that were significantly different from the condition-pooled quantities or significantly predicted compared to a shuffled distribution (Fig. S1DE, Fig. 3E, Fig. 4G, Fig. 5F, Fig. S4DI, Fig. S6G). In order to aggregate over all data to determine whether condition-specific quantities were significantly different from shuffle or significantly predicted within a session relative to shuffle dynamics, we averaged the condition-specific quantity over the relevant dimensions (command, condition, and/or neuron) to yield a single aggregated value for a session. For example in Fig. 3E *right*, we take the distance between average activity for a (command, condition, neuron) tuple and condition-pooled average activity for a (command, neuron) tuple, and we average this distance over (command, condition) tuples to yield an aggregated value that is used to assess if individual neurons are significant. We correspondingly averaged the shuffle distribution across all relevant dimensions (command, condition, and/or neuron). Together this procedure yielded a single aggregated value that could be compared to a single aggregated distribution to determine session significance. Finally, when we sought to aggregate over sessions, we took the condition-specific quantity that was aggregated within a session and averaged it across sessions and again compared it to a shuffle distribution of this value aggregated over sessions.

When R^2^ was the metric assessed (Fig. 4CF, Fig. 5C-E, Fig. S4BFG), a single R^2^ metric was computed for each session and compared to the R^2^ distribution from shuffle models. This R^2^ metric is known as the “coefficient of determination,” and we note that it assesses how well the dynamics-predicted values (e.g. spike counts) account for the variance of the true values.

In some cases, a linear regression was fit between two quantities (Fig. S6CDGJK) on both individual sessions and on data pooled over all sessions, and the significance of the fit and correlation coefficient were both reported. In other cases where random effects such as session or analyzed command may have influenced the linear regression parameters (Fig. S6FG), a Linear Mixed Effect (LME) model was used with session and/or command modeled as random effects on intercept.

In Fig. S5, a paired Student’s t-test was used to compare two models’ R^2^ metric across sessions.Fig. 6 analyzed simulations of OFC models, not experimentally-recorded data. Fig. 6CD used a paired Wilcoxon test and a LME to compare input magnitude between a pair of OFC models. Fig. 6GH used a Mann-Whitney U test and a LME to compare population distance between an OFC model and its shuffle distribution.

