## Supplemental figures for "Invariant neural dynamics drive commands to control different movements"

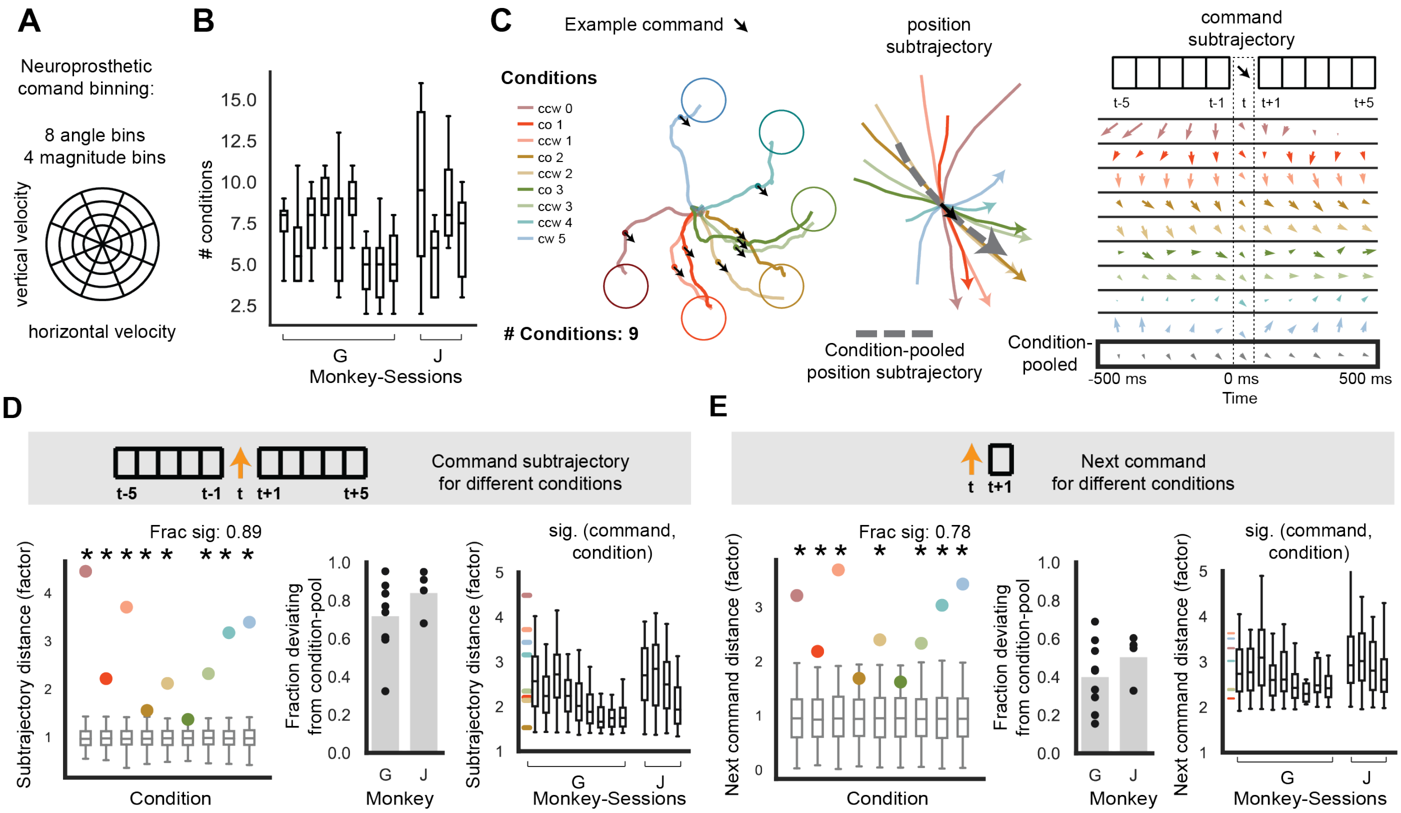


**Figure S1. The same command is issued within different command trajectories to produce different movements. Related to Fig. 1EF.**

**(A)** To analyze the same command in different movements, the continuous-valued two-dimensional commands are categorized into one of 32 bins. Bins discretize the command angle into 8 equally spaced bins and magnitude into 4 bins, so that the pair of angle bin and magnitude bin results in 32 total bins. **(B)** The number of conditions in which the same command occurs frequently enough to be analyzed (>=15 occurrences) for each session analyzed from monkeys G and J. **(C)** *Left*. Repeated from Fig. 1E for visualization: observations of an example command (shown as a black arrow) are plotted during single trials for nine conditions. The example command was in the -45 degree direction and the smallest magnitude bin of analysis (see STAR methods – "Command discretization for analysis"). *Center.* Local cursor position subtrajectory plot (aligned to command occurrence) repeated from Fig. 1F for visualization, plus the condition-pooled cursor position subtrajectory (dashed gray arrow; average over command observations pooling over conditions). *Right.* Command subtrajectory plot repeated from Fig. 1F for visualization, plus the condition-pooled command subtrajectory (gray; average over command observations pooling over conditions). **(D)** Analysis of whether the same command is used within different command subtrajectories in different conditions. The "condition-specific subtrajectory distance" is quantified between each condition-specific command subtrajectory and the condition-pooled command subtrajectory. *Left.* Colored dots show the condition-specific subtrajectory distance for the example command and conditions. The gray boxplots (whiskers span $0^{th}$-$95^{th}$ percentile) show the chance distribution of distances derived from bootstrapping, i.e. subsampling and averaging command subtrajectories from the condition-pooled distribution of command occurrences. For visualization, condition-specific subtrajectory distances are normalized by the mean of the bootstrapped distribution. 89% of the example conditions have command subtrajectories that are significantly different from the condition-pooled command subtrajectory. *Center.* Fraction of (command, condition) tuples with condition-specific command subtrajectories that are significantly different from the condition-pooled command subtrajectory. Condition-specific command subtrajectories are overall significantly different from the condition-pooled command subtrajectory: Monkey G [J]: p-value < 0.001 for 9/9 [4/4] sessions, p-value < 0.001 pooled over sessions (mean of command subtrajectory distances = 1.920 [2.342], mean (95th percentile) of bootstrapped distribution distances = 1.0 (1.01) [1.0 (1.02)]). *Right.* Distribution of condition-specific subtrajectory distances for individually significant (command, condition) tuples. Horizontal colored lines correspond to example conditions shown in left. **(E)**

Analysis of whether the same command is followed by distinct next commands in different conditions. For a given command and condition, the “condition-specific next command” is calculated as the average command following the given command in the given condition. For a given command, the “condition-pooled next command” is the average command following the given command, pooling over conditions. The “condition-specific next command distance” is calculated between the condition-specific next command and the condition-pooled next command. Distances are normalized by the mean of the bootstrapped shuffle distribution. *Left.* Colored dots show the condition-specific next command distance for the example command and conditions. *Center.* Fraction of (command, condition) tuples with condition-specific next commands that are significantly different from the condition-pooled next command. Condition-specific next commands are overall significantly different from the condition-pooled next command: Monkey G [J]: p-value < 0.01 for 9/9 [4/4] sessions, p-value < 0.001 for 8/9 [4/4] sessions, p-value < 0.001 pooled over sessions (mean of next command distances = 1.676 [2.178], mean (95th percentile) of bootstrapped distribution of next command distances = 1.0 (1.03) [1.0 (1.05)]). *Right.* Distribution of condition-specific next command distances for individually significant (command, condition) tuples. Horizontal colored lines correspond to example shown in left.

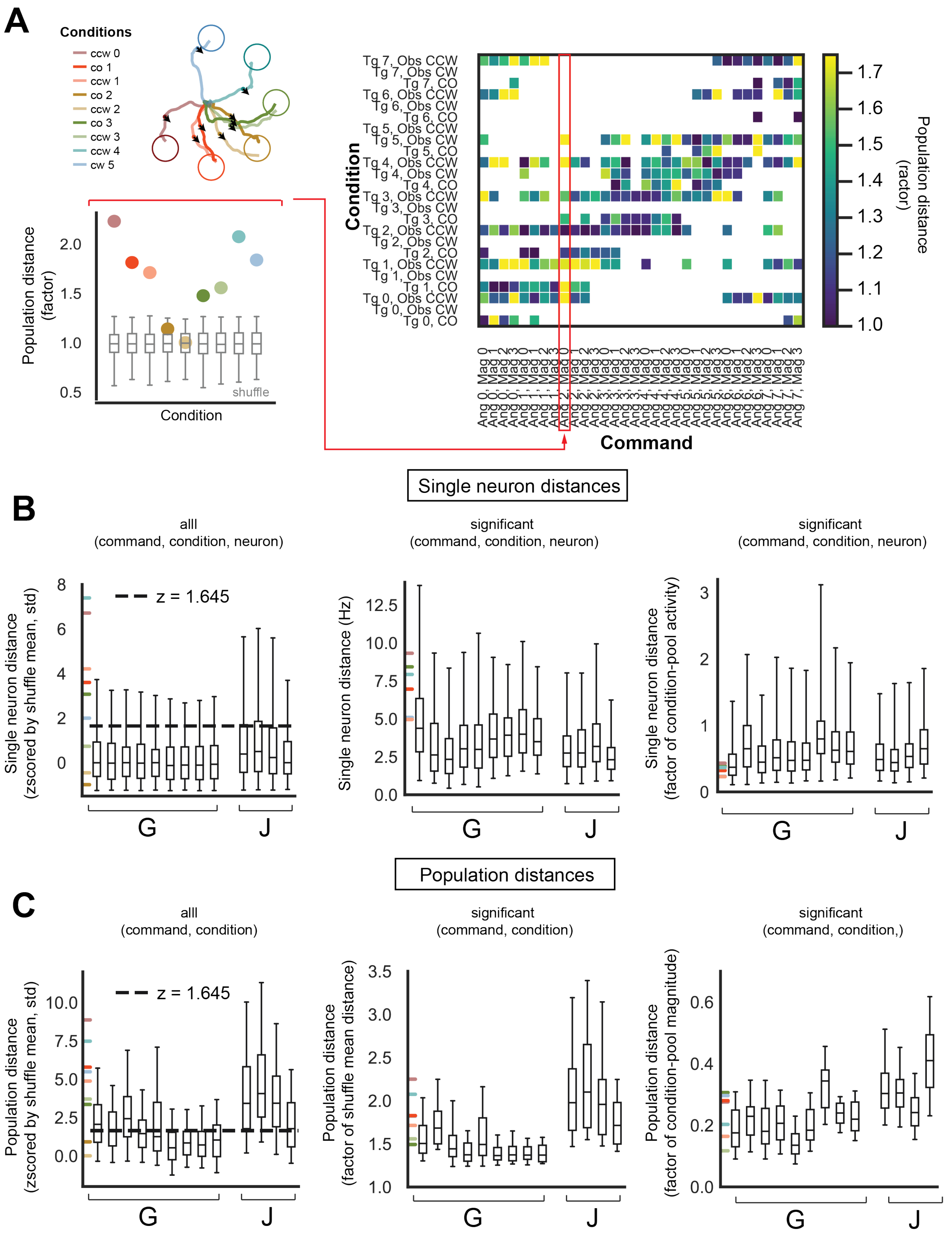

A
Conditions
ccw 0
co 1
ccw 1
co 2
ccw 2
co 3
ccw 3
ccw 4
cw 5
Population distance (factor)
shuffle
Condition
Command
Population distance (factor)
Single neuron distances
B
alll (command, condition, neuron)
significant (command, condition, neuron)
significant (command, condition, neuron)
z = 1.645
Single neuron distance (zscored by shuffle mean, std)
Single neuron distance (Hz)
Single neuron distance (factor of condition-pool activity)
G
J
Population distances
C
alll (command, condition)
significant (command, condition)
significant (command, condition,)
Population distance (zscored by shuffle mean, std)
Population distance (factor of shuffle mean distance)
Population distance (factor of condition-pool magnitude)

**Figure S2. Distributions of condition-specific neural activity issuing a given command. Related to Fig. 3B-E**. In all plots, colored horizontal lines correspond to the example data in Fig. 3B, and whiskers span the 2.5$^{th}$ – 97.5$^{th}$ percentiles of the data distribution.

**(A)** Population activity distances for all (command, condition) tuples in an example session. *Left.* Population activity distances (divided by the shuffle mean) for the example command, conditions, and session (Monkey G, session 0) from Fig. 3B. *Right.* For the same example session, population distances (divided by shuffle mean) for all commands and conditions with sufficient datapoints (>= 15 observations per session) to be analyzed. Columns correspond to the analysis of one command across various conditions (rows). Boxes are not filled in (white) if there are not enough observations of the command in the condition. The example command in *left* is marked in *right* with a red box and the command bin label of "Ang 2, Mag 0". **(B)** Single neuron activity distances. *Left.* For all (command, condition, neuron) tuples, the distance (absolute difference) between condition-specific activity and condition-pooled activity, z-scored by the mean and standard deviation of the shuffle distribution's same distances. The horizontal black line illustrates an estimate of the significance threshold (z = 1.645, 95$^{th}$ percentile of a standard normal distribution). In formal analysis, the empirical (command, condition, neuron) shuffle distribution's 95$^{th}$ percentile serves as the significance threshold for each (command, condition, neuron) tuple. *Center.* For (command, condition, neuron) tuples that are significantly different than shuffle, the distribution of the distance (absolute difference) between condition-specific activity and condition-pooled activity. *Right.* For (command, condition, neuron) tuples that are significantly different than shuffle, the distribution of the distance (absolute difference) between condition-specific activity and condition-pooled activity, divided by the condition-pooled activity. **(C)** Population activity distances. *Left.* For all (command, condition) tuples, the distance between condition-specific activity and condition-pooled activity, z-scored by the mean and standard deviation of the shuffle distribution's same distances. The horizontal black line illustrates an estimate of the significance threshold (z = 1.645, 95$^{th}$ percentile of a standard normal distribution). In formal analysis, the empirical (command, condition) shuffle distribution's 95$^{th}$ percentile serves as the significance threshold for each (command, condition) tuple. *Center.* For (command, condition) tuples that are significantly different than shuffle, the distribution of the distance between condition-specific activity and condition-pooled activity, divided by the mean of the shuffle distribution of the same distance. *Right.* For (command, condition) tuples that are significantly different than shuffle, the distribution of the distance between condition-specific activity and condition-pooled activity, divided by the magnitude of the condition-pooled activity.

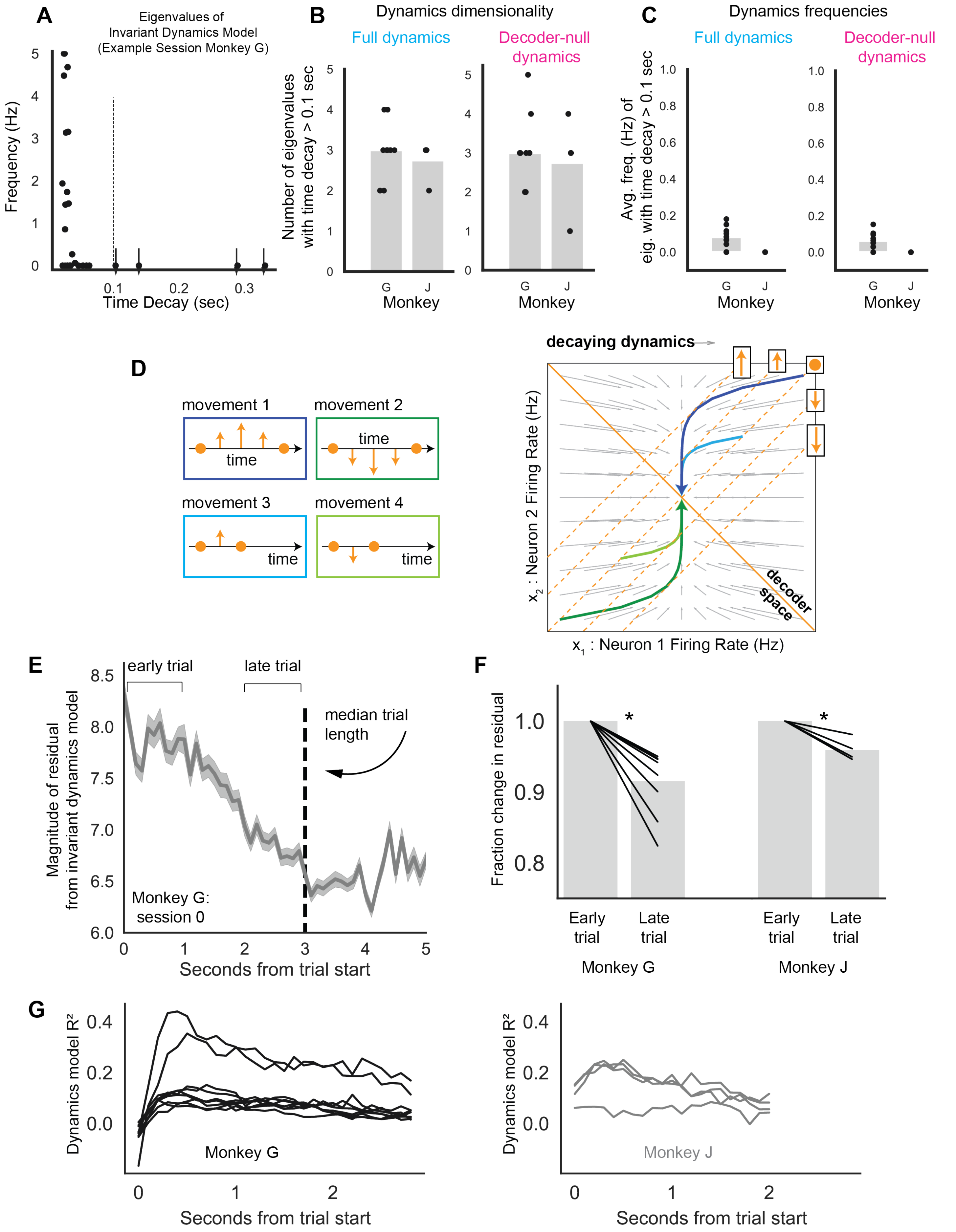

A
Eigenvalues of
Invariant Dynamics Model
(Example Session Monkey G)
Frequency (Hz)
Time Decay (sec)
B
Dynamics dimensionality
Full dynamics
Decoder-null
dynamics
Number of eigenvalues
with time decay > 0.1 sec
Monkey
C
Dynamics frequencies
Full dynamics
Decoder-null
dynamics
Avg. freq. (Hz) of
eig. with time decay > 0.1 sec
Monkey
D
movement 1
movement 2
movement 3
movement 4
time
decaying dynamics
decoder
space
x1 : Neuron 1 Firing Rate (Hz)
x2 : Neuron 2 Firing Rate (Hz)
E
early trial
late trial
median trial
length
Magnitude of residual
from invariant dynamics model
Monkey G:
session 0
Seconds from trial start
F
Fraction change in residual
Early
trial
Late
trial
Monkey G
Monkey J
G
Dynamics model R²
Monkey G
Monkey J
Seconds from trial start

**Figure S3. Properties of invariant dynamics models. Related to Fig. 2, 4C, 6**

**(A)** The frequency and decay properties of the eigenvalues of the dynamics matrix for an example session's dynamics model (Monkey G, session 0). Eigenvalues with decay timescales greater than 0.1 seconds (the timescale at which the BMI updates the command and cursor) are denoted with arrows. **(B)** The dimensionality of the full dynamics matrix (decoder-null dynamics matrix) ranged from 2-4 (2-5) when considering the eigenvalues that had timescales greater than the BMI update's timescale of 0.1 seconds. The number of eigenvalues (each corresponding to one dimension of neural activity) with time decay > 0.1 seconds is shown for each monkey and session. **(C)** The average frequency of the full dynamics matrix (decoder-null dynamics matrix) eigenvalues with time decays > 0.1 seconds is ~0.1 (~0.06) Hz for Monkey G and is 0 Hz (0 Hz) for Monkey J. The average frequency (averaged over eigenvalues with time decays > 0.1 seconds) is shown for each monkey and session. **(D)** *Left.* Example movements which are composed of the same commands in different temporal orders. *Right.* Illustration of neural trajectories that follow invariant, decaying dynamics to control different movements. As illustrated in Fig. 2B, the projection of neural activity into the decoder space determines the command that is issued for movement. As in Fig. 4A, different neural activity patterns are used to issue the same command. For example, the first neural activity pattern in each trajectory is different, although they issue the same command. **(E)** Plot of the magnitude of neural activity that is not explained by the invariant dynamics model (i.e. the residual of the invariant dynamics model's predictions, which is the difference between observed neural activity and the prediction of neural activity based on the previous time step's neural activity). The trial-averaged L2-norm of the residual (divided by the square root of the number of neurons) is shown across time for an example session (Monkey G, session 0). **(F)** The residual magnitude in the late trial period, normalized to the residual magnitude in the early trial period. For analysis, the trial length was set to the median trial time for each session, and the early trial period (late trial period) was the first (last) one-third of the trial length. Each data point is the average of a single session, and the bar is the average across sessions. The residual is larger in the early trial period than the late trial period. Analysis was done using a linear-mixed effect model with session modeled as a random effect and early vs. late modeled as a fixed effect: Monkey G, slope = -0.242, t(4746) = -24.926, p-value = 3.87 $\times 10^{-137}$, Monkey J slope = -0.0625, t(1161) = -5.354, p-value = 8.58 $\times 10^{-8}$. Individual datapoints in the statistics were the average of the norm of the residuals during the early and late epochs for individual trials. Trials that were shorter than the trial median were not included in the statistics. **(G)** The $R^2$ (coefficient of determination) of the invariant dynamics model at each time point relative to the start of the trial, calculated for each session with held-out test data pooling across trials and conditions. The following is some interpretation of this data. At the very start of the trial, the model predicts spiking activity less well, consistent with large input driving neural activity. Then, there is a bump of high predictability, consistent with the initial large input evolving according to invariant dynamics. Then, the predictability decreases to an asymptote, consistent with ongoing feedback modulating neural activity.

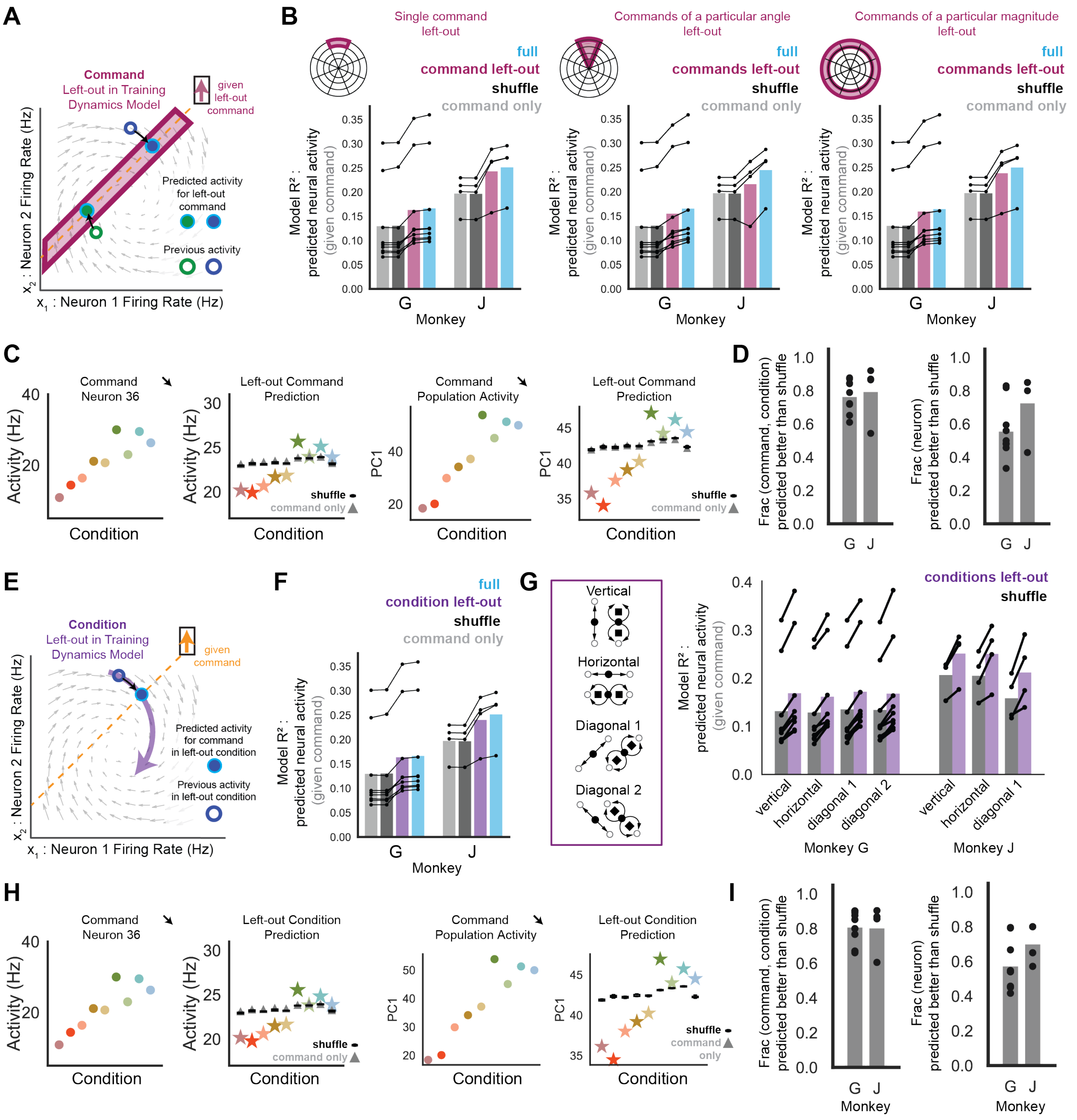


**Figure S4. Generalization of invariant dynamics across sets of commands and conditions. Related to Fig. 4B-E, G.**

As in Fig. 4C, each panel shows the $R^2$ of models predicting neural activity given the command it issues (Monkey G [J]: n=9 [4] sessions) including 1) the dynamics model that is trained using a complete dataset and that predicts held-out test data (cyan, labeled “full”), 2) the dynamics model that is trained with commands left out and that predicts the left-out commands (magenta, labeled “command left-out”), 3) the dynamics model that is trained using a shuffled complete dataset and that predicts held-out, unshuffled test data (black, labeled “shuffle”), and 4) a model trained using a complete dataset and that predicts held-out test data just given the command but not given previous neural activity (gray, labeled “command only”). See STAR methods – “Invariant dynamics models” – “*Generalization of invariant dynamics*”. **(A)** Schematic (as in Fig. 4B *left*). We ask if a linear model of invariant dynamics can predict the neural activity that issues a given command that was left out of training the model. Magenta box indicates that neural activity that transitions to and from the given command are left-out of the dynamics model training data. **(B)** Generalization of invariant dynamics' predictions to sets of commands that were not used to train the invariant dynamics model. Predictions of left-out neural activity are significantly better than shuffle dynamics. *Left.* An individual command is left out and significantly predicted relative to shuffle dynamics (stats reported in Fig. 4C for "command left-out dynamics"). The left-out model coefficient of determination ($R^2$) aggregates the predictions for each left-out command. *Middle.* All commands in a particular angular bin are left out and predicted significantly better than shuffle dynamics (Monkey G [J]: p-value < 0.001 for 9/9 [3/4], p-value n.s for 0/9 [1/4] sessions, p-value < 0.001 for sessions pooled, mean $R^2$ = 0.155 [0.216], mean (95th percentile) $R^2$ of shuffle = 0.130 (0.130) [0.196 (0.196)]). The left-out model $R^2$ aggregates the predictions for each left-out angle of commands. *Right.* All commands in a particular magnitude bin are left out and predicted significantly better than shuffle dynamics (Monkey G [J]: p-value < 0.001 for 9/9 [4/4] sessions, p-value < 0.001 for sessions pooled, mean $R^2$ = 0.159 [0.238], mean (95th percentile) $R^2$ of shuffle = 0.130 (0.130) [0.196 (0.196)]). The left-out model $R^2$ aggregates the predictions for each left-out magnitude of commands. **(C)** Visualization of observed and predicted neural activity for the example command and conditions from Fig. 4DE. Predictions are from the invariant dynamics model that has been trained with data left out for the example command ("left-out command dynamics"). *Left.* Condition-specific average activity for the example neuron and command (repeated from Fig. 4D *left* for visualization). *Left-center.* Prediction for the condition-specific average activity for the example neuron by the left-out dynamics model (stars), the shuffle dynamics model (black boxplot distribution), and the model predicting neural activity only using the command (gray triangle). *Right-center.* Condition-specific average population activity is visualized along the activity dimension that captured the most neural activity variance (the first principal component, labeled "PC1", from principal components analysis applied to condition-specific average population activity) for the example command and conditions (repeated from Fig. 4E *left* for visualization). *Right.* Prediction for the condition-specific average population activity on PC1 by the left-out dynamics model (stars), the shuffle dynamics model (black boxplot distribution), and the model predicting neural activity only using the command (gray triangle). **(D)** Analyses of how well neural activity is predicted for individual (command, condition) tuples when the command is left out of training data for the dynamics model ("left-out dynamics"). *Left.* Fraction of (command, condition) tuples where left-out dynamics predicts condition-specific average population activity significantly better than shuffle dynamics (Monkey G [J]: n=9 [4] sessions). *Right.* Fraction of neurons, aggregated over all (command, condition) tuples, where left-out dynamics predicts the neuron's average activity significantly better than shuffle dynamics (Monkey G [J]: n=9 [4] sessions). **(E)** Schematic (as in Fig. 4B *right*). We ask if the invariant dynamics model can predict neural activity for a given command and condition if all neural activity in that condition (illustrated in purple) is left-out of training the model. **(F)** Predictions of neural activity for a given command in a left-out condition are significantly better than shuffle dynamics (stats reported in Fig. 4C for "condition left-out dynamics"). The left-out model $R^2$ aggregates the predictions for each left-out condition. **(G)** Generalization of invariant dynamics' predictions to sets of conditions that were not used to train the invariant dynamics model. *Left.* Schematics illustrate which conditions were left out and then predicted for each left-out set of conditions. *Right.* All neural activity in a particular set of left-out conditions is left out and predicted significantly better than shuffle dynamics. *Vertical conditions left out*. Monkey G [J]: p-value < 0.001 for 9/9 [4/4] sessions, p-value < 0.001 for sessions pooled (mean $R^2$ = 0.169 [0.251], mean (95th percentile) of shuffled $R^2$ = 0.131 (0.131) [0.206 (0.206)]). *Horizontal conditions left out.* Monkey G [J]: p-value < 0.001 for 9/9 [4/4] sessions, p-value < 0.001 for sessions pooled (mean $R^2$ = 0.161 [0.251], mean (95th percentile) of shuffled $R^2$ = 0.128 (0.128), [0.204, (0.205)]). *Diagonal 1 conditions left out*. Monkey G [J]: p-value < 0.001 for 9/9 [4/4] sessions, p-value < 0.001 for sessions pooled (mean $R^2$ = 0.172 [0.212], mean (95th percentile) of shuffled $R^2$ = 0.135 (0.135) [0.158 (0.158)]). *Diagonal 2 conditions left out*. Monkey G: p-value < 0.001 for 9/9 sessions, p-value < 0.001 for sessions pooled (mean $R^2$ = 0.168, mean (95th percentile) of shuffled $R^2$ = 0.134 (0. 134)). Monkey J: obstacle task did not have these diagonal conditions. **(H)** Visualization of observed and predicted neural activity for the example command and conditions from Fig. 4DE. Predictions are from the invariant dynamics model that has been trained with data left out for each condition separately ("left-out condition dynamics"). Thus, each prediction is made by a separate model with the corresponding condition left out of training data. *Left.* Condition-specific average activity for the example neuron and command (repeated from Fig. 4D *left* for visualization). *Left-center.* Prediction for the condition-specific average activity for the example neuron by the left-out dynamics models (stars), the shuffle dynamics model (black boxplot distribution), and the model predicting neural activity only using the command (gray triangle). *Right-center.* Condition-specific average population activity is visualized along PC1 (see legend (C) for explanation) for the example command and conditions (repeated from Fig. 4E *left* for visualization). *Right.* Prediction for the condition-specific average population activity on PC1 by the left-out dynamics models (stars), the shuffle dynamics model (black boxplot distribution), and the model predicting neural activity only using the command (gray triangle). **(I)** Analyses of how well neural activity is predicted for individual commands and conditions when the condition is left out of training data for the dynamics model ("left-out dynamics"). *Left.* Fraction of (command, condition) tuples where left-out dynamics predicts condition-specific average population activity significantly better than shuffle dynamics (Monkey G [J]: n=9 [4] sessions). *Right.* Fraction of neurons, aggregated over all (command, condition) tuples, where left-out dynamics predicts the neuron's average activity significantly better than shuffle dynamics (Monkey G [J]: n=9 [4] sessions).

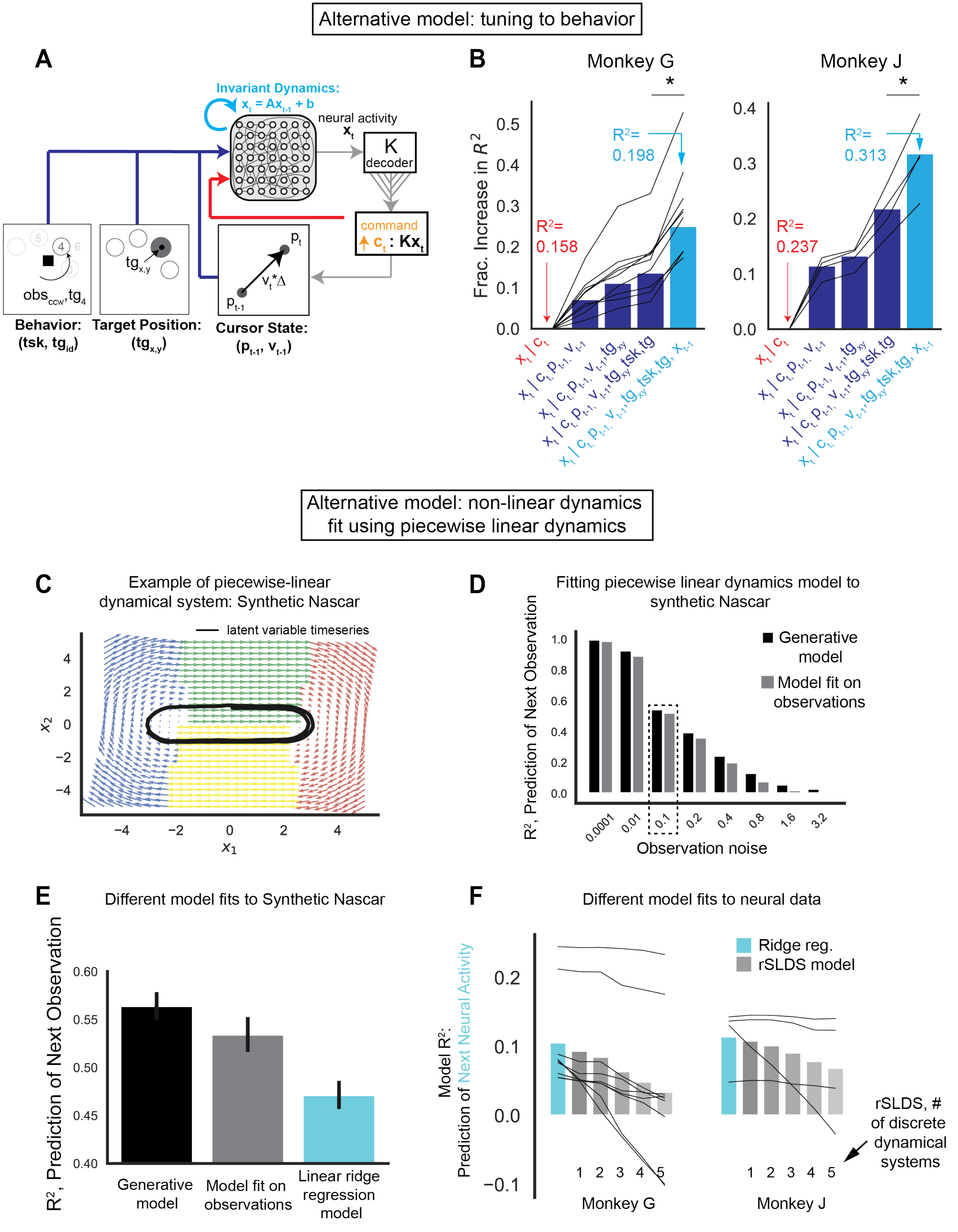
Alternative model: tuning to behavior
A
Invariant Dynamics:
x_t = Ax_t-1 + b
neural activity
x_t
K
decoder
command
c_t : Kx_t
p_t
v_t*Δ
p_t-1
obs_ccw, tg_4
tg_x,y
Behavior:
(tsk, tg_id)
Target Position:
(tg_x,y)
Cursor State:
(p_t-1, v_t-1)
B
Monkey G
Monkey J
Frac. Increase in R^2
R^2= 0.158
R^2= 0.198
R^2= 0.237
R^2= 0.313
x_t | c_t
x_t | c_t, p_t-1, v_t-1
x_t | c_t, p_t-1, v_t-1, tg_xy
x_t | c_t, p_t-1, v_t-1, tg_xy, tsk, tg
x_t | c_t, p_t-1, v_t-1, tg_xy, tsk, tg, x_t-1
Alternative model: non-linear dynamics
fit using piecewise linear dynamics
C
Example of piecewise-linear
dynamical system: Synthetic Nascar
latent variable timeseries
x_1
x_2
D
Fitting piecewise linear dynamics model to
synthetic Nascar
Generative model
Model fit on observations
R^2, Prediction of Next Observation
Observation noise
0.0001
0.01
0.1
0.2
0.4
0.8
1.6
3.2
E
Different model fits to Synthetic Nascar
R^2, Prediction of Next Observation
Generative model
Model fit on observations
Linear ridge regression model
F
Different model fits to neural data
Ridge reg.
rSLDS model
Model R^2:
Prediction of Next Neural Activity
rSLDS, # of discrete dynamical systems
Monkey G
Monkey J

**Figure S5. Alternative models to predict neural activity. Related to Fig. 4AC, 5C.**
Invariant dynamics predict neural activity beyond encoding of cursor, target, and task.
**(A)** Schematic of task-relevant behavior variables that may be encoded in motor cortex population activity $x_t$ at time t during BMI performance, including the command $c_t$ (orange), cursor position $p_{t-1}$ and velocity $v_{t-1}$, target position $tg_{x,y}$ in the 2D workspace, a categorical variable $tg$ that encodes the target, and a categorical variable $tsk$ that encodes task (center-out versus obstacle-avoidance) and whether the trajectory went clockwise vs counterclockwise for the obstacle-avoidance task. **(B)** Fraction increase in coefficient of determination ($R^2$) for models predicting neural activity for a given command $c_t$ as an increasing number of predictors are incorporated. Reported $R^2$ values are on left-out test data, so increasing the number of predictors does not trivially increase $R^2$. Incorporating invariant neural dynamics (cyan) significantly improves upon predictions from the model with all task-relevant variables (right-most dark blue bar "behavior encoding model": $x_t | c_t, p_{t-1}, v_{t-1}, tg_{xy}, tsk, tg$) (Monkey G: paired Student's t-test: N = 18, T = -7.182, p-value = 9.41e-05, Monkey J: paired Student's t-test: N = 8, T = -5.141, p-value = 0.0143).
Non-linear invariant dynamics do not predict neural activity beyond linear invariant dynamics. To test if neural activity predictions may be improved by using a non-linear model of invariant dynamics, we chose to use a recurrent switching linear dynamics system (rsLDS) model [s1]. The rsLDS has the advantage of capturing non-linear dynamics yet still having parameters that are interpretable using linear systems analysis. Specifically, we used a "recurrent-only" switching linear dynamical system that switches between dynamical systems depending only on the latent state [s1]. This was selected for interpretability (i.e. dynamics always obey specific linear dynamics $A$ when the latent state is in a specific region of state space).
**(C)** We first ensured we could properly fit the non-linear dynamics to a toy example, the "Nascar example" copied from [s1] ("ssm" repository -- https://github.com/lindermanlab/ssm) that has activity evolving under a piecewise combination of four linear dynamical systems. **(D)** Forward prediction accuracy of the true Nascar example generative models (black bars) and the fit models (gray bars) confirmed that our fitting procedure found model parameters that yielded comparable accuracy in forward model prediction to the generative model, even when noise was added to observations. Both models suffered similarly from additive noise to the observations. **(E)** In the case of mild additive noise (noise = 0.1, indicated in dotted box in (D)), both the nascar generative (black) and fit (gray) rsLDS models outperformed linear ridge regression (cyan) in prediction of future observations, as expected due to the non-linear generative model. **(F)** Comparison of rsLDS models (gray) vs. linear ridge regression (cyan) fit on neural data as animals perform BMI. We set the latent state dimensionality to the number of neurons that were recorded. This choice was made after sweeping latent state dimensionalities and observing increasing log-likelihoods with higher latent state dimensionality on held-out test data. The linear ridge regression models outperformed the rsLDS models on held-out test data, and the rsLDS performance worsened as more dynamical systems were incorporated (i.e., as more non-linearities were added).

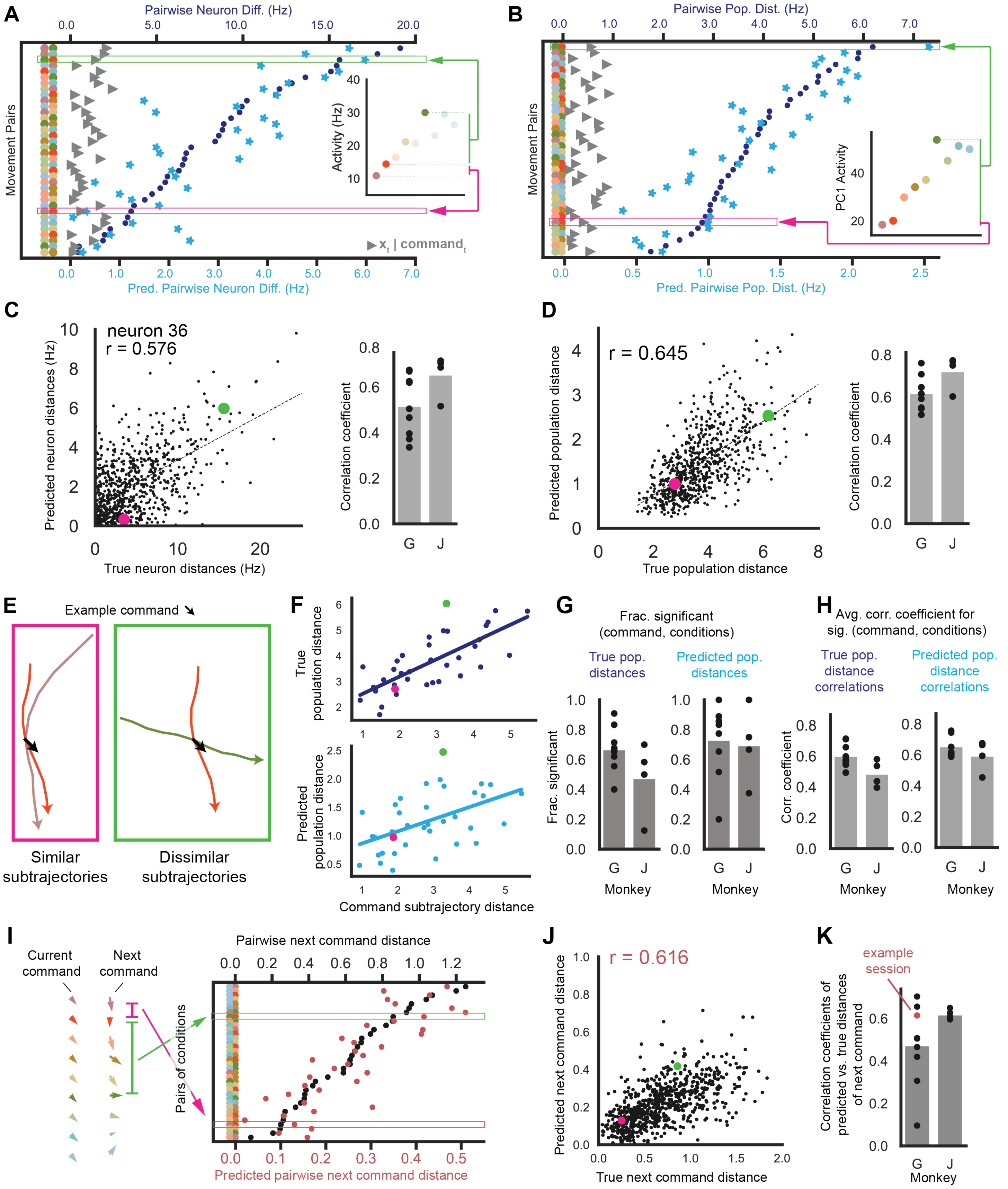


**Figure S6. Invariant dynamics predicts structure across pairs of conditions. Related to Fig. 3, 4ADEG.** **(A)** For a given example command, visualization of the distance between single neuron activity across pairs of conditions (dark blue dots, top x-axis) and the distance between dynamics-predictions of activity across pairs of conditions ("predicted distance", cyan blue dots, bottom x-axis), shown for the command, neuron (neuron 36), and conditions in Fig. 3B. As comparison, activity was predicted given just the continuous-valued commands for each condition in the pair (i.e. activity predicted without an invariant dynamics model), and the distance between these predictions across pairs of conditions is shown (gray triangles which are measured on the cyan scale). The individual conditions composing each condition-pair are indicated with colored dots at the left of the plot. Condition-pairs are sorted by increasing distance. Two examples of condition-pairs are highlighted in pink and green, and the corresponding activity of the individual conditions is shown in the inset. The position subtrajectories for these condition-pairs are shown below in (E). **(B)** Same as (A) but for distances between population activity. Inset shows differences in PC1 activity for illustration, but population distances are calculated in the high-dimensional decoder-null space (see STAR methods – "Analysis of activity issuing a given command"). **(C)** *Left.* For a given command, correlation between the true distance across condition-pairs (x-axis) and predicted distance (y-axis), for the example neuron and session in (A) and Fig. 3B. Dots include all commands and corresponding condition-pairs analyzed for this example session. Pink and green dots indicate condition pairs highlighted in (A). *Right.* Correlation coefficients between true distance across condition-pairs and predicted distance. Each data point is one session, averaging over neurons for all (command, condition-pair) tuples, and the bar is the session-average. Dynamics-predicted distances across condition-pairs are significantly correlated with true distances: Monkey G: p-value < 0.001 for 9/9 sessions, p-value < 0.001 pooled over sessions (linear regression: slope = 0.221, rv = 0.584, N=377489). Monkey J: p-value < 0.001 for 4/4 sessions, p-value < 0.001 pooled over sessions (linear regression: slope = 0.317, rv = 0.693, N = 34643). **(D)** *Left.* Same as (C) *left*, except for population activity distances. *Right.* Same as (C) *right*, except for population activity distances. Each data point is for one session, averaging over all (command, condition-pair) tuples, and the bar is the session-average. Dynamics-predicted distances across condition-pairs are significantly correlated with true distances: Monkey G: p-value < 0.001 for 9/9 sessions, p-value < 0.001 pooled over sessions (linear regression, slope = 0.350, rv = 0.638, N = 5112). Monkey J: p-value < 0.001 for 4/4/ sessions, p-value < 0.001 pooled over sessions (linear regression, slope = 0.449, rv = 0.800, N=1714). **(E)** Analysis of whether neural activity for a given command is more similar across conditions that have more similar command subtrajectories, and whether this structure is predicted by a model of invariant dynamics. Illustration of example condition-pairs used in (A)-(D) with similar (pink) and dissimilar (green) command subtrajectories (position subtrajectories are plotted). **(F)** *Top.* For the example command from Fig. 3B, correlation between population activity distances and command subtrajectory distances for condition-pairs in (A). Pink and green points correspond to pairwise comparisons illustrated in (E). *Bottom.* Same as *Top* but correlation between dynamics-predicted distances and command subtrajectory distances. **(G)** *Left.* Fraction of commands that occur frequently (in > 5 conditions) that exhibit a significant correlation between 1) population activity distance across a condition-pair and 2) command subtrajectory distance across a condition-pair. Over all commands, command subtrajectory distance across a condition-pair is significantly correlated with population activity distance: Monkey G: p-value < 0.001 for 9/9 sessions, pooled over sessions: linear mixed effect (LME) model with command identity and session as random effects: N=4674, z = 31.04, p = 1.36e-211, Monkey J: p-value < 0.05 for 4/4 sessions, p-value < 0.001 for 3/4 sessions, pooled over sessions: LME model with command identity and session as random effects: N=1629, z = 14.18, p-value = 1.22e-45. *Right.* Fraction of commands that occur frequently (in > 5 conditions) that exhibit a significant correlation between 1) the dynamics-predicted population activity distance across a condition-pair and 2) the command subtrajectory distance. Over all commands, command subtrajectory distance across a condition-pair is significantly correlated with dynamics-predicted population activity distance (Monkey G: p-value < 0.01 for 9/9 session, p-value < 0.001 for 8/9 sessions, p-value < 0.001 for pooled sessions (LME with command identity and session modeled as random effects, N = 4674, z = 33.1, p-value = 3.92e-240), Monkey J: p-value < 0.001 for 4/4 sessions, p-value < 0.001 for pooled sessions (LME with command identity and session modeled as random effects, N = 1629, z = 22.5, p-value = 5.93e-112)). **(H)** *Left.* Average correlation coefficient of 1) true population distance across a condition-pair versus 2) command subtrajectory distance, aggregated across significant command-conditions. *Right.* Same as *Left* but correlation of 1) predicted population distance across a condition-pair versus 2) command subtrajectory distance. **(I)** Analysis of how condition-specific neural activity issuing the same current command ($c_t$) transitions forward to issue distinct next commands ($c_{t+1}$). *Left.* For the example command in Fig. 3B, visualization of the average current and next command for each example condition. *Right.* Visualization of the distance between the dynamics-predicted next commands for each condition in a pair (red, bottom x-axis) and the distance between the true next commands across a condition-pair (black, top x-axis). The individual conditions composing each condition-pair are indicated with colored dots at the left of the plot. Condition-pairs are sorted by increasing distance in next command. Two examples of condition pairs are highlighted in pink and green (same as in (A)). **(J)** For the example session, correlation of 1) the distance between true next commands across a condition-pair (x-axis) and 2) the distance between predicted next commands across a condition-pair (y-axis). Dots include all commands and corresponding condition-pairs. Pink and green dots indicate condition-pairs highlighted in (I). **(K)** Same as (J) except for all sessions (example session is shown in red). Dynamics-predicted distance between next commands across a condition-pair is significantly correlated with the true distance: Monkey G: p-value $< 0.001$ for 8/9 sessions, p-value n.s. for 1/9 sessions, p-value $< 0.001$ for pooled sessions (linear regression, slope=0.263, rv =0.76), Monkey J: p-value $< 0.001$ for 4/4 sessions, p-value $< 0.001$ for pooled sessions (linear regression, slope=0.178, rv =0.63).

s1. Linderman, S., Johnson, M., Miller, A., Adams, R., Blei, D., and Paninski, L. (2017). Bayesian Learning and Inference in Recurrent Switching Linear Dynamical Systems. In Proceedings of the 20th International Conference on Artificial Intelligence and Statistics Proceedings of Machine Learning Research., A. Singh and J. Zhu, eds. (PMLR), pp. 914–922.
