## Supplemental tables for "Invariant neural dynamics drive commands to control different movements"

| Analysis description | Figure | Monkey | Session significance | | | Pooled session statistics vs. shuffle | | | |
|---|---|---|---|---|---|---|---|---|---|
| | | | *# sessions with p-value < 0.001* | *# sessions with 0.001 < p-value < 0.05* | *# sessions with p-value > 0.05 (n.s.)* | *mean of data* | *$5^{th}$ percentile of shuffle* | *mean of shuffle* | *$95^{th}$ percentile of shuffle* |
| Distances aggregating over (command, condition, neuron) tuple | 3B | G | 9/9 | | | 1.167 | | 1.004 | 1.010 |
| | | J | 4/4 | | | 1.235 | | 0.745 | 0.757 |
| Population distance aggregating over (command, condition) tuple | 3D | G | 9/9 | | | 1.222 | | 1.0 | 1.007 |
| | | J | 4/4 | | | 1.724 | | 1.0 | 1.019 |
| R2 of full dynamics predictions of neural activity given command (cyan bar) | 4C | G | 9/9 | | | 0.167 | | 0.130 | 0.130 |
| | | J | 4/4 | | | 0.252 | | 0.196 | 0.196 |
| R2 of command left-out dynamics predictions of neural activity given command (magenta bar) | 4C | G | 9/9 | | | 0.163 | | 0.130 | 0.130 |
| | | J | 4/4 | | | 0.243 | | 0.196 | 0.196 |
| R2 of condition left-out dynamics predictions of neural activity given command (purple bar) | 4C | G | 9/9 | | | 0.163 | | 0.130 | 0.130 |
| | | J | 4/4 | | | 0.240 | | 0.196 | 0.196 |
| Single neuron error between true and dynamics-predicted neural activity given command, aggregating over all (command, condition, neuron) tuples | 4D | G | 9/9 | | | 1.232 | 1.359 | 1.359 | |
| | | J | 4/4 | | | 1.182 | 1.454 | 1.455 | |
| Population error between true and dynamics-predicted neural activity given command, aggregating over all (command, condition) tuples, normalized by the mean of the shuffle distribution | 4E | G | 9/9 | | | 0.883 | 0.99 | 1.0 | |
| | | J | 4/4 | | | 0.809 | 0.99 | 1.0 | |
| R2 of condition-specific component of neural activity predicted by dynamics | 4F | G | 9/9 | | | 0.226 | | -0.006 | -0.005 |
| | | J | 4/4 | | | 0.330 | | -0.016 | -0.014 |
| R2 of full dynamics predictions of neural activity (cyan bar) | 5C | G | 9/9 | | | 0.100 | | 0.055 | 0.055 |
| | | J | 4/4 | | | 0.117 | | 0.051 | 0.053 |
| R2 of command left-out dynamics predictions of neural activity (magenta bar) | 5C | G | 9/9 | | | 0.099 | | 0.055 | 0.055 |
| | | J | 4/4 | | | 0.113 | | 0.051 | 0.053 |
| R2 of condition left-out dynamics predictions of neural activity (purple bar) | 5C | G | 9/9 | | | 0.097 | | 0.055 | 0.055 |
| | | J | 4/4 | | | 0.103 | | 0.051 | 0.053 |
| R2 of decoder-null dynamics predictions of neural activity (pink bar) | 5C | G | 9/9 | | | 0.083 | | 0.055 | 0.055 |
| | | J | 4/4 | | | 0.085 | | 0.051 | 0.053 |
| R2 of full dynamics predictions of command (orange bar) | 5D | G | 9/9 | | | 0.315 | | 0.264 | 0.266 |
| | | J | 4/4 | | | 0.212 | | 0.186 | 0.188 |
| R2 of command left-out dynamics predictions of command (magenta bar) | 5D | G | 9/9 | | | 0.310 | | 0.264 | 0.266 |
| | | J | 4/4 | | | 0.211 | | 0.186 | 0.188 |
| R2 of condition left-out dynamics predictions of command (purple bar) | 5D | G | 9/9 | | | 0.305 | | 0.264 | 0.266 |
| | | J | 2/4 | 1/4 | 1/4 | 0.193 | | 0.186 | 0.188 |
| R2 of decoder-null dynamics predictions of command (pink bar) | 5D | G | | | 9/9 | 0.0 | | 0.264 | 0.266 |
| | | J | | | 4/4 | 0.0 | | 0.186 | 0.188 |
| Error in prediction of condition-specific next command | 5E | G | 9/9 | | | 3.956 | 5.38 | 5.40 | |
| | | J | 4/4 | | | 7.324 | 9.240 | 9.305 | |
| Fraction of (command, condition) tuples with sign of next command's angle is accurately predicted with full dynamics | 5G | G | 9/9 | | | 0.708 | | 0.535 | 0.541 |
| | | J | 4/4 | | | 0.617 | | 0.473 | 0.480 |
| Error in prediction of next command's angle with full dynamics | 5G | G | 9/9 | | | 19.503 | 26.415 | 26.564 | |
| | | J | 4/4 | | | 9.608 | 12.636 | 12.779 | |

**Table S1. Comparisons to shuffled datasets. Related to Figures 3-5**. Statistics computed for individual animal sessions and pooling across sessions compared to shuffled datasets as described in main text and STAR Methods.

| Analysis description | Figure | Monkey | Session significance | | Pooled session statistics (linear mixed effect model with session modeled as random effect) | | |
|---|---|---|---|---|---|---|---|
| | | | *Individual session comparison test* | *# of sessions w/ p-value < 0.05* | *Number of datapoints (N)* | *z statistic* | *p-value* |
| Input magnitude, comparison between full dynamics and no dynamics | 6C | G | Wilcoxon signed-rank test with conditions paired | 9/9 | 432 | 10.49 | 9.67e-26 |
| | | J | | 4/4 | 192 | 5.20 | 1.92e-7 |
| Input magnitude, comparison between decoder-null dynamics and no dynamics | 6D | G | Wilcoxon signed-rank test with conditions paired | 0/9 | 432 | 0.002 | 0.998 |
| | | J | | 0/4 | 192 | -0.003 | 0.990 |
| Distance between average population activity for a (command, condition) compared to shuffle: Full dynamics (cyan) | 6G | G | Mann-Whitney U test | 9/9 | 4906 | -23.09 | 6.37e-118 |
| | | J | | 4/4 | 2408 | -16.68 | 1.77e-62 |
| Distance between average population activity for a (command, condition) compared to shuffle: No dynamics (black) | 6G | G | Mann-Whitney U test | 0/9 | 4334 | 0.168 | 0.866 |
| | | J | | 0/4 | 2188 | 0.462 | 0.644 |
| Distance between average population activity for a (command, condition) compared to shuffle: Decoder-null dynamics (pink) | 6H | G | Mann-Whitney U test | 0/9 | 4488 | 0.932 | 0.351 |
| | | J | | 0/4 | 2252 | -1.490 | 0.136 |
| Distance between average population activity for a (command, condition) compared to shuffle: No dynamics (black) | 6H | G | Mann-Whitney U test | 0/9 | 4482 | 0.611 | 0.541 |
| | | J | | 0/4 | 2250 | 0.449 | 0.654 |

**Table S2. Simulation statistics. Related to Figure 6**. Statistics computed for simulations performed on individual animal sessions and pooling across sessions as described in main text and STAR Methods.
